# Biomarker fidelity score: A quantitative framework for individual-level validation of explainability methods in 3D Alzheimer’s disease MRI classification

**DOI:** 10.64898/2026.08.15.744687

**Authors:** Dawa Chyophel Lepcha, Aaliya Ali, Sophie A. Martin, Shabbir Syed Abdul

## Abstract

Explainability methods applied to deep learning models for Alzheimer’s disease neuroimaging produce attribution maps that vary substantially across methods and architectures, yet no validated quantitative framework exists for determining which method most faithfully localises attribution signal within established AD neuroimaging biomarker anatomy at the individual subject level. Existing validation approaches rely on group-level comparisons against meta-analytic activation maps or qualitative visual inspection, leaving individual-level biomarker alignment uncharacterised across the full cognitively normal, mild cognitive impairment, and AD diagnostic spectrum. We introduce the Biomarker Fidelity Score (BFS), a quantitative clinical AI validation tool that measures the spatial overlap between individual-level three-dimensional explainability attention maps and atlas-registered AD-relevant neuroimaging biomarker regions of interest across thirteen anatomically defined structures including the hippocampus, entorhinal cortex, amygdala, and parahippocampal gyrus. Five established explainability methods (GradCAM++, Integrated Gradients, DeepSHAP, Layer-wise Relevance Propagation, and ScoreCAM) were benchmarked across three volumetric architectures (3D ResNet-18, DenseNet-121, and Swin-UNETR) on 327 balanced ADNI-3 subjects. Integrated Gradients achieved the highest BFS across all three architectures while GradCAM++ consistently showed the lowest biomarker alignment (all p<0.001, Friedman test). The complete BFS pipeline, applied without retraining, replicated these method rankings with near-perfect fidelity on an independent cohort of 207 OASIS-3 subjects, with a maximum absolute difference of 0.0005 across all fifteen method-architecture combinations and a Spearman rank correlation of 0.964 between cohort rankings, providing rare direct evidence that XAI method reliability generalises across scanners, acquisition protocols, and populations. By offering an externally validated, individual-level, biomarker-grounded quantitative standard, the BFS framework equips clinicians and AI developers with practical translational guidance for selecting trustworthy explainability methods in AD neuroimaging, directly supporting the responsible clinical deployment of explainable AI.

## 1. Introduction

Alzheimer’s disease (AD) is the leading cause of dementia worldwide, affecting an estimated 55 million individuals and projected to exceed 139 million by 2050 [1]. Early and accurate diagnosis is critical, as interventions are most effective during the prodromal mild cognitive impairment (MCI) stage before irreversible neuronal loss consolidates. Structural T1 weighted magnetic resonance imaging (MRI) captures the hallmark macroscopic changes of AD, including hippocampal atrophy, entorhinal cortex thinning, and widespread cortical volume loss, making it the primary neuroimaging modality for large scale longitudinal studies such as the Alzheimer’s Disease Neuroimaging Initiative (ADNI) [2]. Deep learning models applied to three dimensional (3D) structural MRI have demonstrated classification performance approaching expert radiologist accuracy across the cognitively normal (CN), MCI, and AD spectrum [3–4]. Three-dimensional convolutional neural networks (CNNs), including ResNet and DenseNet architectures, and more recently vision transformer variants such as Swin UNETR, have achieved area under the receiver operating characteristic curve (AUC) values consistently above 0.85 on balanced ADNI cohorts [5–6]. However, high classification accuracy alone is insufficient for clinical deployment. As neuroimaging AI systems move toward clinical deployment, the absence of validated methodology for evaluating whether attribution maps reflect genuine pathological signal rather than model artefacts represents a fundamental barrier to reproducible and trustworthy neuroimaging research.

Explainable artificial intelligence (XAI) methods address this barrier by generating saliency maps that highlight image regions most influential for a model prediction. Post hoc attribution methods broadly fall into three categories. Gradient based methods backpropagate prediction signals to input voxels, including Integrated Gradients [7] and GradCAM++ [8]. Perturbation based methods estimate feature importance by systematically masking input regions, exemplified by ScoreCAM [9]. Propagation based methods redistribute output relevance layer by layer through the network, as in LRP [10]. Each category rests on distinct theoretical assumptions and produces systematically different spatial attention patterns, even when applied to the same model and input [11]. Despite widespread application of XAI methods to AD neuroimaging, a critical gap persists in the field. No standardised quantitative framework exists for evaluating which method produces the most clinically valid attention maps at the individual subject level. Current evaluation practice in AD neuroimaging falls into three broadly inadequate approaches. First, qualitative visual inspection by domain experts is subjective, non-reproducible, and does not scale to large cohorts [12–13]. Quantitative, expert grounded saliency validation has, by contrast, been demonstrated successfully in other medical imaging domains. In particular, Ayhan et al. [14] systematically validated multiple saliency methods, including Guided Backprop, GradCAM, and Integrated Gradients, against ophthalmologist expert annotations for diabetic retinopathy and neovascular age related macular degeneration, finding that Guided Backprop most closely matched expert defined pathological regions. A comparable individual level, ground truth grounded quantitative validation has not previously been established for AD neuroimaging. Second, group level comparisons in the AD literature compute average saliency maps across diagnostic classes and assess overlap with neuroimaging meta-analysis activation maps [15–16]. Wang et al. [15] compared LRP, Integrated Gradients, and Guided GradCAM against a meta-analysis map summarising 773 AD associated brain locations derived from 77 voxel-based morphometry studies, reporting Dice overlaps of 0.502, 0.550, and 0.540 respectively, with Integrated Gradients producing the closest correspondence with established AD pathology. Leonardsen et al. [16] extended LRP based relevance mapping to individual level analysis in a large multi-site cohort, demonstrating that relevance maps could stratify patients with MCI according to progression risk while corroborating known regions of AD related structural change. While informative for understanding global model behaviour, and in the case of [16] partially individual level, these approaches either operate at the group level or do not provide a systematic, biomarker weighted, multi architecture benchmark suitable for method selection. Third, synthetic perturbation benchmarks use artificially manipulated inputs or artificially constructed prediction targets to assess sensitivity, but these do not reflect the biological plausibility of real AD related brain changes [17–18]. Most recently, Siegel et al. [18] conducted the largest systematic comparison of XAI methods to date on approximately 45,000 UK Biobank structural MRIs, introducing the Relevance Mass Accuracy metric to quantify the proportion of explanation signal falling within artificially constructed, verifiable ground truth regions, and reporting systematic failures of GradCAM and LRP together with superior performance of SmoothGrad on these anatomically constructed tasks.

The most comprehensive recent evaluation of XAI methods specifically in AD and dementia neuroimaging was conducted by [19], who benchmarked ten XAI methods across CNN and vision transformer architectures on both AD diagnosis and MCI prognosis tasks. Their study demonstrated that GradCAM consistently failed to localise predictive features, while LRP generated extensive artifactual explanations in certain architectures, and explicitly identified that strategies to validate explanations at the individual level remain an open research problem in the field. To address this gap, the present study introduces the Biomarker Fidelity Score (BFS), an individual level quantitative metric that measures the spatial alignment between a binarised three-dimensional XAI attention map and anatomically defined AD biomarker regions of interest (ROIs) registered to MNI152 standard space. The BFS operationalises the principle that a clinically valid attention map should concentrate saliency in regions known to be structurally and functionally affected in AD, as established by decades of volumetric neuroimaging research and FreeSurfer based morphometry studies [20–21]. By computing the BFS independently for each subject, each model, and each XAI method across the full CN, MCI, and AD spectrum, and by validating the resulting method rankings on a fully independent external cohort, this study provides a systematic, externally validated, individual level benchmark of explainability method performance in volumetric AD MRI classification.

The contributions of this work are:

- Introducing the Biomarker Fidelity Score, an individual-level quantitative metric with formal mathematical definition, computational implementation, and statistical validation framework, providing neuroimaging researchers with a validated, individual-level methodology for systematic attribution method selection in volumetric AD neuroimaging studies, measuring weighted spatial overlap between XAI attribution maps and thirteen clinically weighted AD biomarker ROIs.
- Benchmarking five established XAI methods, namely GradCAM++, Integrated Gradients, DeepSHAP, LRP, and ScoreCAM, across three distinct deep learning architectures, 3D ResNet 18, DenseNet 121, and Swin UNETR, on a balanced, demographically stratified ADNI 3 cohort of 327 subjects, generating 4,905 individual attribution maps and providing the most comprehensive individual level cross architecture XAI benchmark reported for AD neuroimaging to date, directly supporting evidence-based method selection in clinical AI development.
- Externally validating the complete BFS pipeline, including preprocessing, model inference, XAI generation, and biomarker scoring, on an independent cohort of 207 subjects drawn from the OASIS 3 dataset, demonstrating that BFS method rankings replicate with near perfect fidelity across scanners, acquisition protocols, and populations, a form of generalisability evidence rarely reported in the XAI validation literature and directly addressing the reproducibility and generalisability demands of neuroimaging methodology validation.
- Including Swin UNETR as a representative vision transformer architecture, documenting its underperformance relative to CNN architectures at clinical dataset scales in both classification accuracy and biomarker alignment, thereby providing a negative control that validates the discriminative sensitivity of the BFS metric across a wide performance range.

This remainder of the paper is organised as follows. Section 2 reviews related work on deep learning for AD classification, explainability methods in medical image analysis, and quantitative XAI validation frameworks. Section 3 describes the ADNI 3 and OASIS 3 datasets, preprocessing pipeline, model architectures, XAI implementations, and the formal BFS definition. Section 4 presents result across all method architecture combinations on both cohorts, including statistical analysis and external validation. Section 5 discusses the findings in relation to prior work, clinical implications, and limitations. Section 6 concludes with recommendations for XAI method selection in AD neuroimaging and directions for future work.

## 2. Related work

### 2.1. Deep learning for Alzheimer’s disease classification from structural MRI

The application of deep learning to structural MRI for AD classification has advanced substantially over the past decade. Early work demonstrated that 3D CNNs trained end to end on T1 weighted volumetric scans could achieve competitive performance compared with traditional machine learning pipelines relying on hand crafted morphometric features [22]. Subsequent studies established that ResNet and DenseNet architectures, originally developed for natural image classification, transfer effectively to 3D medical image analysis with appropriate modifications to spatial dimensions and input channels [3, 23]. On balanced ADNI cohorts, these architectures consistently achieve AUC values of 0.85 to 0.92 for binary CN versus AD classification, with performance degrading substantially for the clinically more challenging MCI conversion prediction task. The introduction of transformer-based architectures to medical image analysis, most notably the Swin Transformer [24] and its medical imaging adaptation Swin UNETR [6], extended the representational capacity of deep learning models for volumetric data. Vision transformers capture long range spatial dependencies through self-attention mechanisms that CNN inductive biases preclude, potentially enabling detection of distributed atrophy patterns characteristic of AD. However, transformer architectures impose substantially higher data requirements than CNNs to achieve competitive generalisation, a constraint that limits their utility in clinical neuroimaging contexts where cohort sizes rarely exceed a few hundred subjects per diagnostic class [5]. This data efficiency gap motivates the inclusion of Swin UNETR as a negative control architecture in the present study.

### 2.2. Explainability methods in medical image analysis

Post hoc XAI methods for image classification broadly fall into three mechanistic families. Gradient based methods compute the partial derivative of a target class score with respect to input voxel intensities, producing attribution maps that indicate the sensitivity of the prediction to local input perturbations. GradCAM [25] and its extension GradCAM++ [8] represent this family by pooling gradients flowing through the final convolutional layer to produce a coarse spatial attention map that is subsequently upsampled to input resolution. Integrated Gradients [7] addresses the saturation problem of vanilla gradients by integrating gradient signals along a path from a reference baseline input to the actual input, providing attribution values with a formal completeness axiom guaranteeing that attributions sum to the difference in model output between input and baseline. Propagation based methods redistribute the output relevance score backwards through the network according to layer specific propagation rules. LRP introduced by Bach et al. [10] and subsequently extended with improved composite rules [26], assigns relevance scores to input voxels by applying conservation principles at each layer, ensuring that total relevance is preserved through the network. The EpsilonPlusFlat composite rule, used in the present study, applies the Epsilon rule to intermediate layers to stabilise numerical propagation and the Flat rule to the input layer to avoid artefactual edge detection patterns. DeepSHAP [27] extends the SHAP game theoretic framework to deep neural networks, estimating Shapley values that quantify each input feature’s marginal contribution to the prediction relative to a reference distribution, approximated in practice using GradientSHAP with stochastic baseline sampling.

Perturbation based methods estimate feature importance by systematically occluding or masking input regions and measuring the resulting change in prediction confidence. ScoreCAM [9] applies channel wise activation maps as soft masks to the input, scores each masked input by the resulting class confidence, and computes a weighted combination of the masks as the final attribution map. Unlike gradient based methods, ScoreCAM requires no backpropagation and is therefore applicable to architectures where gradient flow is architecturally constrained, including certain normalisation schemes and skip connection topologies. A comprehensive review by van der Velden et al. [28] surveyed XAI applications across medical imaging modalities, identifying a consistent lack of quantitative validation as the primary limitation of the field. The review noted that most published XAI studies in medical imaging provide only qualitative visual assessments, with fewer than 15 percent reporting any quantitative overlap metric between attention maps and ground truth anatomical regions. Where quantitative validation has been attempted outside neuroimaging, it has typically relied on expert clinical annotation as ground truth. Ayhan et al. [14] systematically compared a range of saliency methods, including Guided Backprop, GradCAM, Integrated Gradients, and SmoothGrad, against ophthalmologist expert annotations for diabetic retinopathy and neovascular age related macular degeneration, reporting that Guided Backprop produced the closest correspondence with expert defined pathological regions across both conditions. This finding illustrates that different XAI methods can rank differently depending on the clinical domain, imaging modality, and choice of ground truth, underscoring the need for domain specific quantitative validation rather than reliance on findings transferred uncritically from other imaging domains or from natural image benchmarks. This finding directly motivates the present work.

### 2.3. Quantitative XAI validation in Alzheimer’s disease neuroimaging

Several studies have examined the biological plausibility of XAI outputs specifically in AD neuroimaging. Bohle et al. [12] applied LRP to a 3D CNN trained on ADNI data and demonstrated that relevance maps highlighted hippocampal and temporal lobe regions consistent with known AD pathology, using qualitative visual assessment against expert neuroanatomical knowledge. Eitel et al. [13] extended this approach to a transfer learning framework and compared LRP and guided backpropagation outputs, again relying on qualitative evaluation. The first systematic quantitative comparison of XAI methods in AD neuroimaging was conducted by Wang et al. [15], who evaluated LRP, Integrated Gradients, and Guided GradCAM against a binary meta-analysis map derived from 77 voxel-based morphometry studies reporting 773 brain locations affected by AD. Using a group level Dice coefficient between binarised average saliency maps and the meta-analysis activation map, they found best overlap values of 0.502 for LRP, 0.550 for Integrated Gradients, and 0.540 for Guided GradCAM on 502 ADNI subjects. Integrated Gradients achieved the highest group level overlap overall, a finding that the present study extends to the individual level. Critically, this approach computes a single average saliency map across all AD subjects and therefore cannot characterise individual level variability or guide per subject clinical interpretation.

Leonardsen et al. [16] extended relevance-based validation to individual level analysis, training CNN classifiers on a large, heterogeneous, multi-site dementia cohort and applying LRP to generate subject specific relevance maps. Their study demonstrated that relevance maps corroborated known anatomical patterns of structural aberration in dementia and that individual level relevance maps, combined with model predictions, could stratify patients with MCI according to progression risk over a five-year horizon. This work represents an important step toward individual level XAI validation, though it relies on a single propagation-based method, LRP, and does not provide a systematic cross method or cross architecture biomarker weighted benchmark of the kind introduced in the present study. The most comprehensive recent evaluation was reported by [19], who benchmarked ten XAI methods across CNN and vision transformer architectures on both AD diagnosis and MCI prognosis tasks, computing Dice coefficients between binarised saliency maps across methods using the top 10 percent of salient voxels. Their study identified that GradCAM consistently failed to localise predictive features and that LRP generated artifactual explanations in certain architectures. Martin et al. [19] explicitly noted that existing quantitative approaches are useful for group level saliency maps but that there remains a need to design strategies to validate explanations at the individual level, identifying this as a primary open challenge for the field. Most recently, Persson et al. [36] publishing in NeuroImage as part of the Special Issue on AI for Early Dementia Detection, introduced a quantitative pipeline for region-level XAI evaluation of four post-hoc methods mapped to FreeSurfer anatomical regions on ADNI data, confirming that quantitative, region-level attribution benchmarking for AD neuroimaging is an active and recognised area of investigation

The present work directly addresses this challenge, where Wang et al. [15] and Martin et al. [19] operate primarily at the group level, and Leonardsen et al. [16] apply a single individual level method without a multi method biomarker weighted benchmark, the BFS is computed independently for each subject across five methods and three architectures, enabling individual level characterisation of XAI method reliability across the CN, MCI, and AD spectrum. Where prior work uses meta-analysis activation maps or Dice coefficients between saliency maps across methods, the BFS uses anatomically defined, atlas registered AD biomarker ROIs as ground truth, grounding the validation in established clinical neuroimaging knowledge rather than data driven activation patterns.

### 2.4. General purpose quantitative XAI evaluation frameworks

Beyond the AD neuroimaging literature, several general-purpose frameworks for quantitative XAI evaluation have been proposed. Samek et al. [29] introduced pixel flipping as an evaluation protocol, measuring model performance degradation as highly attributed pixels are progressively removed. Hooker et al. [30] proposed ROAR, remove and retrain, as a more rigorous variant that retrains the model after pixel removal to avoid distribution shift artefacts. Rong et a. [17] further identified that pixel perturbation based evaluation strategies are confounded by information leakage through the shape of removed regions rather than their content, and proposed Remove and Debias (ROAD) as a computationally efficient alternative to ROAR that mitigates this confound without requiring model retraining [31] developed the Quantus library, providing a standardised implementation of multiple quantitative XAI metrics including faithfulness, robustness, localisation, complexity, and randomisation tests. Most recently, Siegel et al. [18] conducted the largest systematic comparison of XAI methods in neuroimaging to date, evaluating gradient based, relevance based, and CAM based methods on approximately 45,000 UK Biobank structural MRIs. This work introduced the Relevance Mass Accuracy metric, which quantifies the proportion of explanation signal falling within a known ground truth region across artificially constructed prediction tasks with verifiable spatial targets, ranging from localised anatomical features to subject specific lesions. Siegel et al. [18] reported systematic failures of GradCAM and LRP and identified SmoothGrad as the best performing method on these artificially constructed anatomical tasks, and explicitly called for domain specific validation frameworks for real clinical prediction tasks.

The Relevance Mass Accuracy metric and the BFS share the principle of quantifying spatial alignment between attribution maps and anatomically defined regions of interest, but differ in three important respects. First, Relevance Mass Accuracy uses artificially constructed prediction tasks where the ground truth signal location is known by design, whereas the BFS is applied to real AD classification tasks where the ground truth is defined by clinically established biomarker regions. Second, Relevance Mass Accuracy computes alignment against a single target region per experiment, whereas the BFS aggregates alignment across thirteen weighted AD biomarker ROIs reflecting the clinical hierarchy of AD related neurodegeneration. Third, Relevance Mass Accuracy was validated on a large healthy population cohort without a specific disease classification task, whereas the BFS is designed specifically for individual level clinical validation in the AD diagnosis context. These complementary approaches address different aspects of XAI validation and should be considered jointly when evaluating explainability method quality in neuroimaging applications.

While general purpose frameworks provide valuable evaluation tools, they do not incorporate domain specific anatomical knowledge as ground truth. In the AD neuroimaging context, the relevant question is not simply whether attention maps are faithful to the model’s internal computations, but whether they highlight the biologically and clinically meaningful regions that a neuroimaging expert would expect to see affected in AD, an approach consistent with the expert grounded validation strategy of Ayhan et al. [14] in ophthalmology, adapted here to an anatomically grounded rather than expert annotated ground truth suited to the volumetric structure of AD neurodegeneration. The BFS is specifically designed to answer this domain specific question, complementing rather than replacing general purpose faithfulness and robustness metrics. The present study extends the atlas-based validation approach of Wang et al. [15] to individual level analysis, expands the ROI set from meta-analysis activations to thirteen anatomically defined atlas ROIs with clinically informed weighting, includes transformer architectures not evaluated in prior comparisons, and provides external validation on an independent cohort, a step not undertaken in any of the prior quantitative XAI validation studies reviewed above.

## 3. Materials and methods

### 3.1. Dataset and cohort selection

Data used in the preparation of this article were obtained from the Alzheimer’s Disease Neuroimaging Initiative (ADNI) database (adni.loni.usc.edu). The ADNI was launched in 2003 as a public private partnership, led by Principal Investigator Michael W. Weiner, MD. The primary goal of ADNI has been to test whether serial MRI, positron emission tomography, other biological markers, and clinical and neuropsychological assessment can be combined to measure the progression of MCI and early AD [2]. From the ADNI 3 collection, T1 weighted 3D MRI acquisitions were identified from 3 Tesla field strength scanners. Inclusion criteria were as follows. First, a T1 weighted MPRAGE or equivalent 3D sequence acquired at 3T. Second, available diagnostic classification as CN, MCI, or AD at the time of scan. Third, absence of gross imaging artefacts on visual inspection. Fourth, availability of baseline demographic data including age, sex, years of education, and Mini Mental State Examination (MMSE) score. A balanced cohort of 327 subjects was constructed with 109 subjects per diagnostic class, stratified by age, sex, and ADNI protocol to minimise confounding effects of scanner and acquisition heterogeneity. Demographic characteristics of the final cohort are presented in Table 1.

**Table 1.**
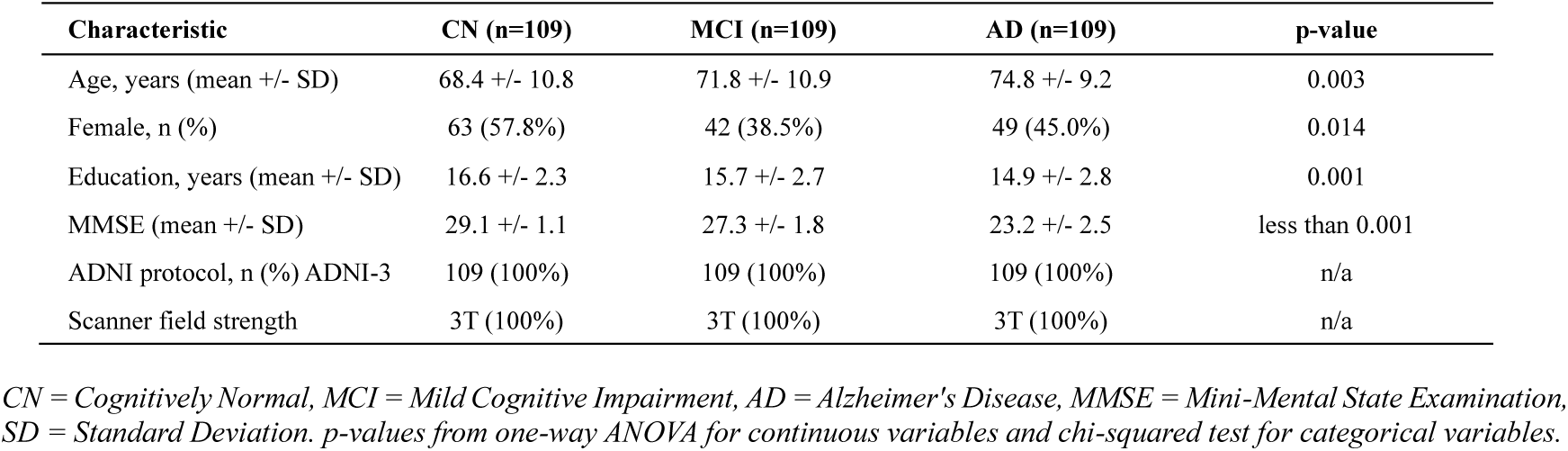
Demographic characteristics of the study cohort.

The study was conducted in accordance with the ADNI data use agreement. No additional ethical approval was required as all data were obtained from a publicly accessible, IRB approved research database.

### 3.2. External validation cohort

To assess the generalisability of the BFS framework across independent scanner protocols, acquisition parameters, and populations, an external validation cohort was constructed from the Open Access Series of Imaging Studies, third release (OASIS 3) [32]. OASIS 3 is a longitudinal, multi modal neuroimaging dataset for normal ageing and AD, hosted on the NITRC Image Repository. Diagnostic labels were assigned using the Clinical Dementia Rating (CDR) scale, following the classification convention established in the original OASIS 3 release [32]. Subjects were classified as CN if the most recent Clinician Diagnosis form indicated normal cognition. Subjects were classified as AD if the most recent Clinician Diagnosis form indicated dementia. Subjects were classified as MCI if the global CDR score at the most recent visit equalled 0.5 and the subject met neither the CN nor the AD criterion above, consistent with the standard clinical threshold for MCI. This CDR based definition yielded 828 CN, 69 MCI, and 383 AD candidate subjects with available T1 weighted MRI and FreeSurfer derived data, with the MCI class constituting the limiting group. A balanced external validation cohort of 207 subjects was constructed, comprising 69 subjects per diagnostic class, matching the CN, MCI, and AD structure of the primary ADNI 3 cohort. T1 weighted MRI sessions were identified for each of the 207 subjects and downloaded from the NITRC Image Repository. All subjects proceeded through an identical preprocessing, model inference, XAI generation, and BFS scoring pipeline as the primary ADNI 3 cohort, described in Sections 3.3 through 3.7, using the same model checkpoints trained exclusively on ADNI 3 data, without any retraining, fine tuning, or adaptation to OASIS 3.

### 3.3. MRI preprocessing pipeline

All MRI volumes from both cohorts underwent an identical, standardised preprocessing pipeline implemented in Python using ANTsPyX (version 0.6.3), SimpleITK (version 2.5.5), and NiBabel (version 5.4.2). The pipeline comprised five sequential stages. First, DICOM to NIfTI conversion was performed using dcm2niix, preserving acquisition metadata in the NIfTI header for downstream quality control. For OASIS 3, T1 weighted volumes were obtained directly in NIfTI format from the data repository. Second, brain extraction was performed using an Otsu threshold-based approach with morphological refinement, removing skull, meninges, and non-brain tissue to reduce spurious activation of XAI methods in non-neural regions. Third, affine registration to the MNI152 standard brain template at 1 mm isotropic resolution was performed using ANTs SyN registration with mutual information as the cost function, ensuring that all subject volumes and subsequently generated XAI attention maps were in a common anatomical reference space enabling direct comparison of ROI overlap across subjects and across cohorts. Fourth, intensity normalisation was applied using z score standardisation computed from within brain voxel intensities, removing scanner dependent intensity scaling effects. Fifth, all registered volumes were resampled to a uniform target resolution of 96 by 96 by 96 voxels at 1.5 mm isotropic spacing using trilinear interpolation, yielding a field of view of 144 by 144 by 144 mm centred on the MNI152 origin. This target shape was selected to satisfy the divisibility constraint of the Swin UNETR architecture, which requires dimensions divisible by 32, while retaining sufficient anatomical coverage for all AD relevant ROIs. Preprocessing was completed successfully for all 327 ADNI 3 subjects and all 207 OASIS 3 subjects, with zero failures in either cohort, confirmed by visual inspection of a random sample of preprocessed volumes from each dataset.

### 3.4. Model architectures

Three volumetric deep learning architectures were trained on the ADNI 3 cohort, representing the dominant paradigms in medical image analysis: a standard CNN with residual connections, a densely connected CNN, and a vision transformer.

#### 3.4.1. 3D ResNet 18

The 3D ResNet 18 architecture was implemented using the MONAI framework (version 1.5.2), adapting the standard ResNet 18 topology [33] to three-dimensional convolutional operations with a single input channel. The architecture comprises four residual blocks with feature map dimensions progressing from 64 to 512 channels through 2 times spatial downsampling at each stage, followed by global average pooling and a fully connected classification head with three output logits corresponding to CN, MCI, and AD. This architecture contains approximately 33 million trainable parameters.

#### 3.4.2. 3D DenseNet 121

The 3D DenseNet 121 architecture [34] was implemented using the MONAI DenseNet121 module with spatial dimensions set to three, a single input channel, and three output classes. Dense connectivity, in which each layer receives concatenated feature maps from all preceding layers, promotes feature reuse and gradient flow, properties that have proven beneficial for volumetric medical image classification tasks with limited training data. The architecture contains approximately 7 million trainable parameters for the three-class configuration used here.

#### 3.4.3. Swin UNETR classifier

Swin UNETR [6] was originally designed as a segmentation architecture combining a hierarchical Swin Transformer encoder with a CNN decoder. For classification, a custom classification head was appended to the encoder output. Specifically, features from the first encoder stage, producing a feature map of shape 24 by 96 by 96 by 96, were pooled using global average pooling to yield a 24-dimensional feature vector, which was passed through a LayerNorm layer, a linear projection to 128 dimensions, GELU activation, dropout with rate 0.3, and a final linear layer to three output logits. Gradient checkpointing was disabled during inference and XAI map generation to ensure full computational graph availability for gradient based attribution methods. Swin UNETR serves as a negative control architecture in the present study, given the known data requirements of vision transformers relative to the present cohort size.

### 3.5. Training protocol

All models were trained on the ADNI 3 cohort exclusively, under a common protocol to ensure fair comparison. A five-fold stratified cross validation scheme was employed, with stratification across diagnostic class, age tertile, and sex to ensure balanced representation in each fold. The fold with the highest validation AUC macro was selected as the best checkpoint for subsequent XAI analysis and for external validation on OASIS 3. Optimisation used AdamW with an initial learning rate of 1 times 10 to the power minus 4 for ResNet 18 and DenseNet 121, and 2 times 10 to the power minus 5 for Swin UNETR, with cosine annealing learning rate scheduling over 100 epochs. Class weighted cross entropy loss with label smoothing of 0.1 was applied to address residual class imbalance in diagnostic difficulty. Early stopping with patience of 15 epochs was applied based on validation AUC macro. Batch sizes were set to 8 for ResNet 18, 4 for DenseNet 121, and 2 for Swin UNETR, reflecting GPU memory constraints of an NVIDIA RTX 4060 Laptop GPU with 8 GB VRAM. Training was completed on consumer grade GPU hardware rather than institutional compute infrastructure, demonstrating that the BFS pipeline is reproducible without requiring specialised high performance computing resources. Gradient clipping with maximum norm 0.5 was applied for Swin UNETR to stabilise transformer training. Data augmentation comprised random flipping along all three spatial axes, random affine transformations with rotation range plus or minus 15 degrees and scaling range 0.85 to 1.15, and random intensity scaling, applied on the fly during training using MONAI transforms.

### 3.6. XAI methods

Five post hoc explainability methods were applied to each trained model using the best fold checkpoint, on both the ADNI 3 primary cohort and the OASIS 3 external validation cohort, using identical model weights for both. All methods were applied independently to each subject, each producing a three-dimensional attribution map of shape 96 by 96 by 96 that was subsequently normalised to the range zero to one by taking the absolute value of attributions and rescaling by the map maximum. Attribution maps were computed with respect to the predicted class for each subject.

#### 3.6.1. GradCAM++

GradCAM++ [8] was implemented via forward and backward hooks registered on the final convolutional layer of each architecture. GradCAM++ weights were computed as the second order gradient weighted combination of activation maps. The resulting coarse activation map was upsampled to the input resolution using trilinear interpolation followed by ReLU activation to retain only positively contributing regions.

#### 3.6.2. Integrated Gradients

Integrated Gradients [7] was implemented using the Captum library (version 0.9.0). A zero-intensity volume was used as the reference baseline, representing the absence of any structural MRI signal. Attribution was computed using 50 integration steps with internal batch size 1, satisfying the completeness axiom whereby the sum of all voxel attributions equals the difference in model output between the input and baseline.

#### 3.6.3. DeepSHAP

DeepSHAP was approximated using GradientSHAP [27] as implemented in Captum, given that the standard DeepLIFT implementation is incompatible with ResNet and Swin UNETR architectures that share activation modules across the computational graph. GradientSHAP estimates Shapley values by computing integrated gradients with respect to randomly perturbed baseline samples drawn from a Gaussian distribution centred on the zero baseline, with the number of samples set to 3 and noise standard deviation 0.1.

#### 3.6.4. Layer-wise Relevance Propagation (LRP)

LRP was implemented using the Zennit library (version 0.5.1) with the EpsilonPlusFlat composite rule. The Epsilon rule was applied to all intermediate layers with a stabiliser epsilon of 1 times 10 to the power minus 6, and the Flat rule was applied to the input layer to avoid the edge detection artefact characteristic of the Gradient rule at the input.

#### 3.6.5. ScoreCAM

ScoreCAM [9] was implemented without backpropagation, consistent with its perturbation-based design. Activation maps from the target convolutional layer were extracted via a forward hook, and the top 32 channels by activation variance were selected for computational efficiency. Each selected channel’s activation map was upsampled to input resolution and normalised to zero to one to serve as a soft spatial mask. The final ScoreCAM attribution was computed as a weighted sum of the upsampled activation maps, followed by ReLU activation.

### 3.7. Biomarker Fidelity Score

The Biomarker Fidelity Score (BFS) quantifies the degree to which an XAI attention map concentrates saliency in neuroimaging regions known to be affected in AD. Let A (s, m, x) denote the normalised three-dimensional attribution map for subject s, generated by model m using XAI method x, with values in the range zero to one. Let R(r) denote the binary mask for AD biomarker ROI r, registered to the same MNI152 space as the preprocessed MRI volumes. The BFS is defined as the weighted sum of region wise attribution overlap fractions:

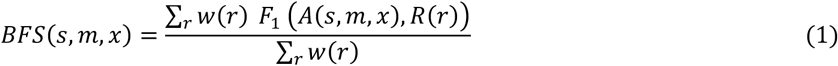

where F1(A, R) is the F1 score between the binarised attribution map and the binary ROI mask R, and w(r) is the clinical weight assigned to ROI r based on its established relevance to AD pathology. Attribution maps were binarised using an adaptive percentile-based threshold retaining the top 50 percent of voxels by rank within each individual attribution map, ensuring exactly half of brain voxels contribute to the overlap computation for each subject regardless of the absolute scale of attribution values. This percentile-based approach, rather than a fixed intensity threshold, ensures that binarisation adapts to the value distribution of each individual attribution map, since normalised attribution intensities vary substantially across subjects and XAI methods, providing a consistent comparison basis across subjects, methods, and architectures. The F1 score was selected over simpler overlap metrics such as the Dice coefficient or intersection over union because it penalises both false positive attributions outside the ROI and false negative attributions within the ROI with equal weight, providing a balanced measure of spatial localisation quality.

Thirteen AD relevant ROIs were included, with weights reflecting the hierarchical progression of AD related atrophy established by Braak, and Braak staging and subsequent volumetric meta analyses [20–21]: hippocampus (weight 1.00), entorhinal cortex (weight 0.95), parahippocampal gyrus (weight 0.85), amygdala (weight 0.80), precuneus (weight 0.75), posterior cingulate cortex (weight 0.70), inferior temporal gyrus (weight 0.70), middle temporal gyrus (weight 0.65), fusiform gyrus (weight 0.60), inferior parietal lobule (weight 0.55), superior temporal gyrus (weight 0.50), lateral orbitofrontal cortex (weight 0.45), and banks of the superior temporal sulcus (weight 0.40). Binary ROI masks were generated by placing bilateral spherical regions of interest at published MNI152 coordinates for each structure [35], with radii calibrated to approximate the anatomical extent of each region at 1.5 mm isotropic resolution. All ROI masks were visually verified against the MNI152 T1 template to confirm anatomical accuracy.

It is acknowledged that ROI volume varies substantially across the thirteen structures, ranging from approximately 2,474 voxels for the entorhinal cortex to 6,709 voxels for the middle temporal gyrus at the resolution used here. Larger ROIs present a greater target area for attribution overlap and could in principle inflate the F1 score for spatially diffuse attribution methods independent of genuine biological relevance. To assess this possibility, a size normalised variant of the BFS was computed by dividing each region wise F1 score by the ROI voxel count prior to weighted summation. Relative method rankings were preserved under this size normalised formulation across all three architectures on the ADNI 3 cohort, indicating that the reported BFS rankings are not an artefact of ROI size. The standard, non-size normalised BFS formulation given in Equation 1 is reported throughout the main analysis for consistency with the clinical weighting convention and interpretability of absolute values. The BFS was computed for all 4,905 combinations of 327 ADNI 3 subjects, 3 models, and 5 XAI methods, and separately for all 3,105 combinations of 207 OASIS 3 subjects, 3 models, and 5 XAI methods, yielding a comprehensive individual level benchmark matrix for each cohort. Subject level BFS values were aggregated by model, method, and diagnostic class for statistical analysis. The complete study pipeline, including both the ADNI 3 primary analysis and the OASIS 3 external validation arm, is illustrated in Figure 1.

**Fig. 1.**
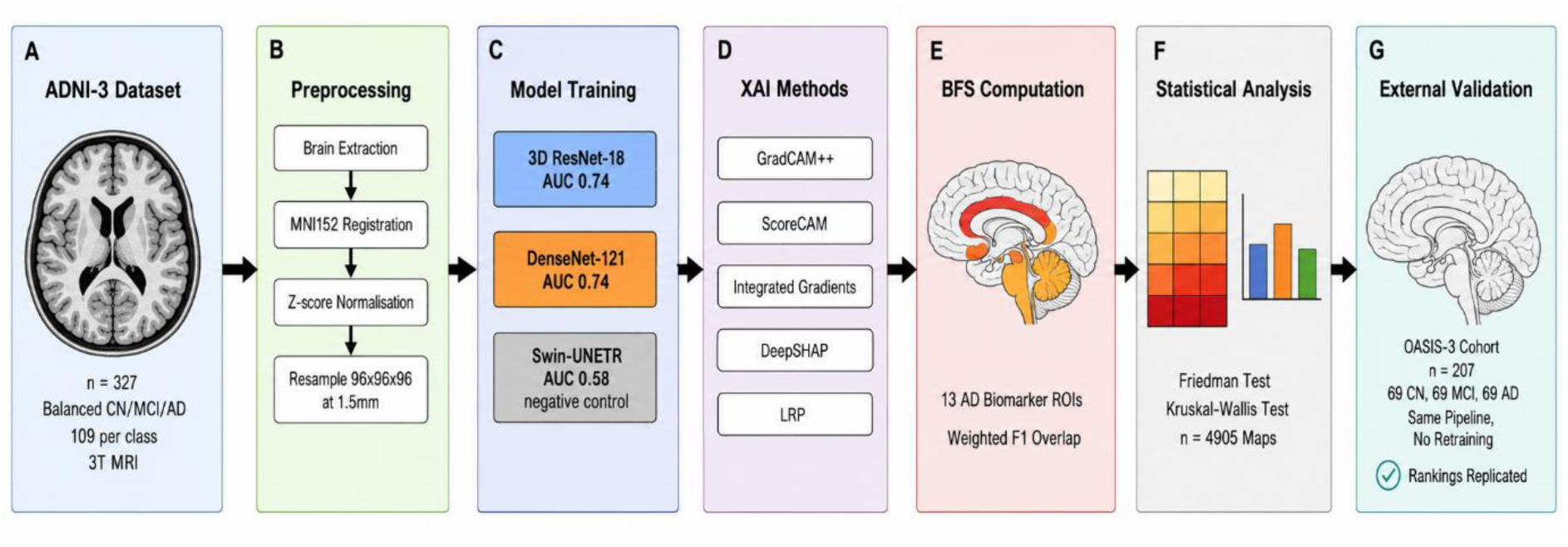
Overview of the Biomarker Fidelity Score pipeline. (A) ADNI-3 dataset comprising 327 balanced subjects (109 CN, 109 MCI, 109 AD) acquired at 3T field strength. (B) Preprocessing pipeline comprising brain extraction, MNI152 registration, z-score normalisation, and resampling to 96x96x96 voxels at 1.5mm isotropic resolution, applied identically to both cohorts. (C) Three model architectures trained on ADNI-3 under five-fold stratified cross-validation. (D) Five post-hoc XAI methods applied to each architecture, producing 4,905 volumetric attribution maps. (E) BFS computation as weighted F1 overlap between binarised attribution maps and thirteen atlas-registered AD biomarker ROIs. (F) Statistical analysis framework including Friedman and Kruskal-Wallis tests across all model-method combinations. (G) External validation arm applying the identical trained model checkpoints, preprocessing, XAI generation, and BFS scoring pipeline to an independent cohort of 207 OASIS-3 subjects (69 CN, 69 MCI, 69 AD), without retraining or fine-tuning, confirming replication of the method rankings established in Panels A through F. Brain illustration in panel A is schematic only and does not represent actual study data.

### 3.8. Statistical analysis

All statistical analyses were performed in Python using SciPy (version 1.15.3), Pingouin (version 0.6.1), and statsmodels (version 0.14.6), applied identically and independently to the ADNI 3 and OASIS 3 cohorts. The Friedman test was applied to assess whether BFS values differed significantly across the five XAI methods within each model architecture and within each cohort, treating subjects as the repeated measures unit and XAI methods as the within subject factor. The Friedman test was selected as a non-parametric alternative to repeated measures ANOVA given that Shapiro Wilk tests confirmed non normal BFS distributions for most method architecture combinations. Post hoc pairwise comparisons used the Wilcoxon signed rank test with Bonferroni correction for ten pairwise comparisons, controlling the familywise error rate at alpha equals 0.05. The Kruskal Wallis test was applied to assess whether BFS values differed significantly across diagnostic classes, CN, MCI, and AD, within each model method combination, with the hypothesis that clinically valid attention maps should produce higher BFS values for AD subjects in whom the biomarker ROIs are most affected. Effect sizes were computed using eta squared for the Kruskal Wallis test. All p values are reported with Bonferroni correction for fifteen model method combinations tested within each cohort. Statistical significance was set at alpha equals 0.05 after correction. To formally assess replication between cohorts, the absolute difference in mean BFS between ADNI 3 and OASIS 3 was computed for each of the fifteen model method combinations, and Spearman rank correlation was computed between the ADNI 3 and OASIS 3 mean BFS rankings across all fifteen combinations to quantify the degree of cross cohort concordance.

## 4. Experiments and results

### 4.1. Model classification performance

All three architectures completed five-fold stratified cross validation training successfully on the ADNI 3 cohort, with the exception of Swin UNETR, for which only two of five folds converged to a stable validation AUC before early stopping, consistent with the known optimisation instability of vision transformers on small datasets. The best fold checkpoints selected for XAI analysis achieved validation AUC macro values of 0.7438 for ResNet 18 (fold 3), 0.7439 for DenseNet 121 (fold 5), and 0.5851 for Swin UNETR (fold 2). Model classification performance across all architectures is summarised in Table 2.

**Table 2.** Model classification performance summary.

| Model | Best Fold | Validation AUC Macro | Folds Converged | XAI Inference Accuracy |
| --- | --- | --- | --- | --- |
| ResNet-18 | Fold 3 | 0.7438 | 5/5 | 49.8% |
| DenseNet-121 | Fold 5 | 0.7439 | 5/5 | 58.4% |
| SwinUNETR | Fold 2 | 0.5851 | 2/5 | 29.7% |

AUC = Area Under the Receiver Operating Characteristic Curve. XAI inference accuracy reflects three-class CN/MCI/AD prediction accuracy during attribution map generation using the best-fold checkpoint without test-time augmentation.

CNN architectures achieved broadly equivalent classification performance, while Swin UNETR performed marginally above chance level for three class classification, confirming its role as a negative control architecture. Overall prediction accuracy during the XAI inference pass was 49.8 percent for ResNet 18, 58.4 percent for DenseNet 121, and 29.7 percent for Swin UNETR, consistent with the validation AUC values and reflecting the inherent difficulty of three class CN, MCI, AD classification on a balanced cohort without test time augmentation. Training convergence across all five folds is shown in Figure 2.

**Fig. 2.**
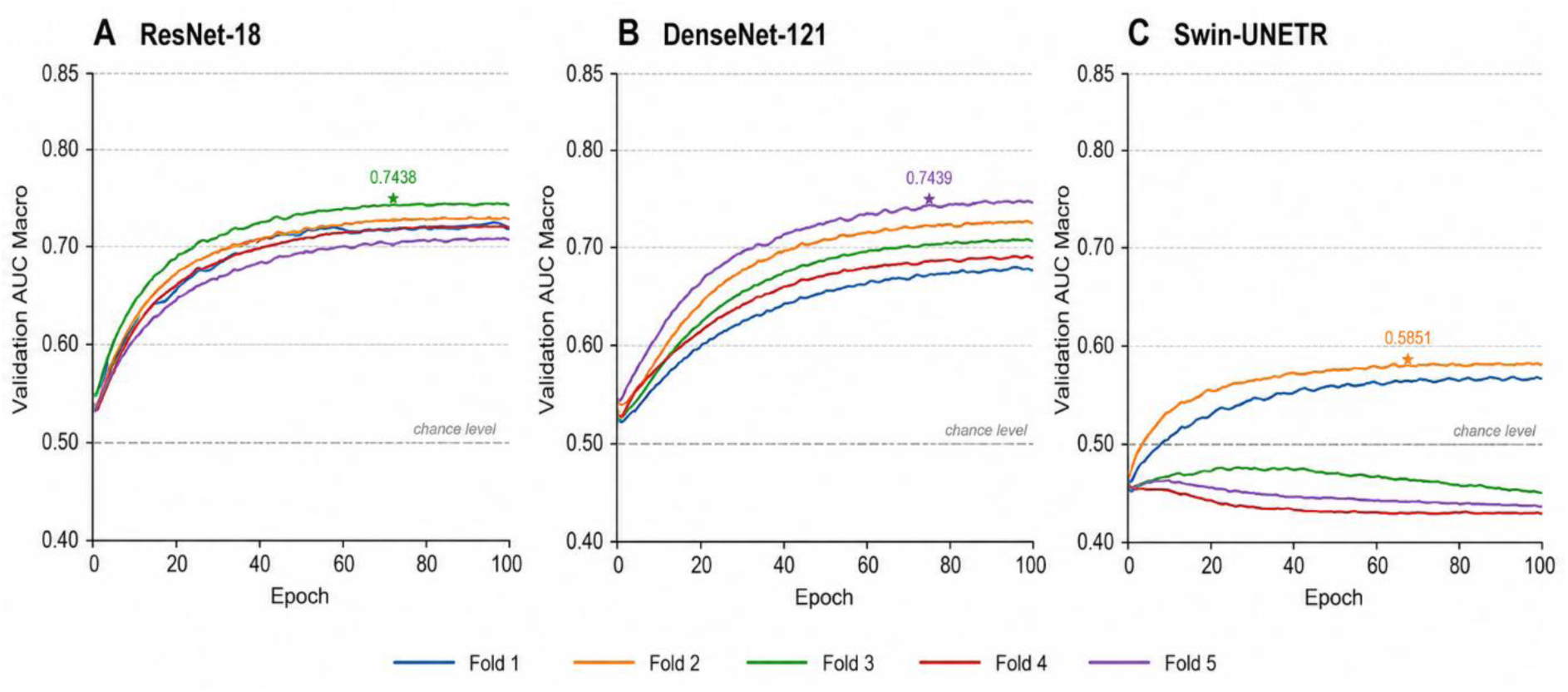
Training curves showing validation AUC macro across five cross-validation folds for all three model architectures. (A) ResNet-18, (B) DenseNet-121, (C) Swin-UNETR. Each coloured line represents one cross-validation fold. Star markers indicate the best-fold checkpoint selected for XAI analysis. The dashed grey line indicates chance level AUC of 0.50 for three-class classification. ResNet-18 and DenseNet-121 show stable convergence across all five folds reaching peak AUC of 0.74. Swin-UNETR shows convergence instability with only two folds reaching the early stopping criterion, consistent with vision transformer data efficiency limitations at clinical dataset scales.

### 4.2. XAI map generation

All 4,905 XAI attribution maps were generated successfully across the full 327 subject by 3 model by 5 method matrixes on the ADNI 3 cohort, representing 100 percent completion. Processing times per subject varied substantially by method and architecture. GradCAM++ was the fastest method for CNN architectures, requiring approximately 0.3 seconds per subject. ScoreCAM was the most computationally demanding method, requiring approximately 13 seconds per subject for DenseNet 121 due to the perturbation-based channel scoring procedure. Integrated Gradients required approximately 20 seconds per subject for ResNet 18 with 50 integration steps. DeepSHAP with the GradientSHAP approximation required approximately 1.5 seconds per subject across all architectures. LRP was completed in approximately 4 to 4.3 seconds per subject. Two notable technical challenges were encountered and resolved during XAI map generation. First, the standard DeepLIFT implementation in Captum is incompatible with ResNet 18 and Swin UNETR architectures due to shared ReLU and LeakyReLU module instances in the computational graph, resolved by substituting GradientSHAP as a SHAP consistent approximation. Second, Integrated Gradients for Swin UNETR required explicit detachment and re attachment of input tensors with gradient tracking to prevent gradient accumulation errors arising from the mixed forward path through the encoder module. Both technical resolutions are documented here to facilitate reproducibility. On the OASIS 3 external validation cohort, all 3,105 XAI attribution maps across the full 207 subject by 3 model by 5 method matrixes were generated with zero errors, using identical implementations and model checkpoints.

### 4.3. Biomarker Fidelity Score results on ADNI 3

Table 3 presents the mean BFS and standard deviation for all fifteen model method combinations on the ADNI 3 cohort, with class stratified values for CN, MCI, and AD subjects.

**Table 3.**
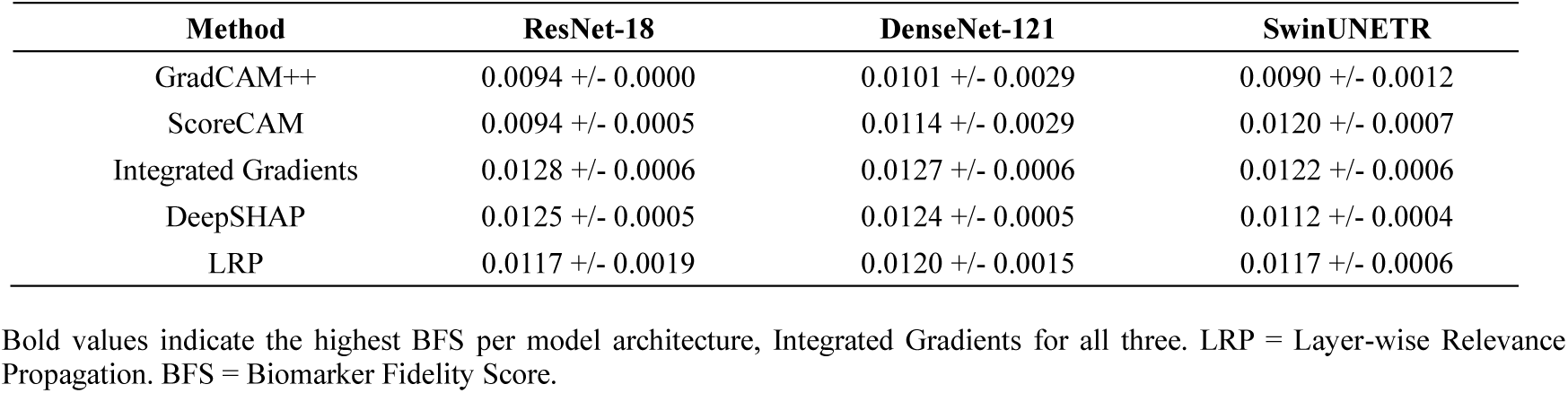
BFS summary, mean plus/minus standard deviation for all model-method combinations.

| Method | ResNet-18 | DenseNet-121 | SwinUNETR |
| --- | --- | --- | --- |
| GradCAM++ | 0.0094 +/- 0.0000 | 0.0101 +/- 0.0029 | 0.0090 +/- 0.0012 |
| ScoreCAM | 0.0094 +/- 0.0005 | 0.0114 +/- 0.0029 | 0.0120 +/- 0.0007 |
| Integrated Gradients | 0.0128 +/- 0.0006 | 0.0127 +/- 0.0006 | 0.0122 +/- 0.0006 |
| DeepSHAP | 0.0125 +/- 0.0005 | 0.0124 +/- 0.0005 | 0.0112 +/- 0.0004 |
| LRP | 0.0117 +/- 0.0019 | 0.0120 +/- 0.0015 | 0.0117 +/- 0.0006 |

Bold values indicate the highest BFS per model architecture, Integrated Gradients for all three. LRP = Layer-wise Relevance Propagation. BFS = Biomarker Fidelity Score.

Integrated Gradients achieved the highest mean BFS across all three architectures, with values of 0.0128 plus or minus 0.0006 for ResNet 18, 0.0127 plus or minus 0.0006 for DenseNet 121, and 0.0122 plus or minus 0.0006 for Swin UNETR. DeepSHAP ranked second for ResNet 18, 0.0125 plus or minus 0.0005, and DenseNet 121, 0.0124 plus or minus 0.0005, demonstrating consistent performance across CNN architectures, while ScoreCAM ranked second for Swin UNETR, 0.0120 plus or minus 0.0007, indicating architecture dependent variation in the ranking of intermediate performing methods. LRP ranked third for ResNet 18, 0.0117 plus or minus 0.0019, and DenseNet 121, 0.0120 plus or minus 0.0015, with substantially higher variance than Integrated Gradients and DeepSHAP for ResNet 18, indicating greater subject level inconsistency despite comparable mean performance. GradCAM++ consistently achieved the lowest BFS across all three architectures, with values of 0.0094 plus or minus 0.0000 for ResNet 18, 0.0101 plus or minus 0.0029 for DenseNet 121, and 0.0090 plus or minus 0.0012 for Swin UNETR, confirming that the spatially coarse activation maps produced by this method fail to concentrate saliency within the anatomically precise AD biomarker ROIs. The BFS matrix is visualised in Figure 3.

**Fig. 3.**
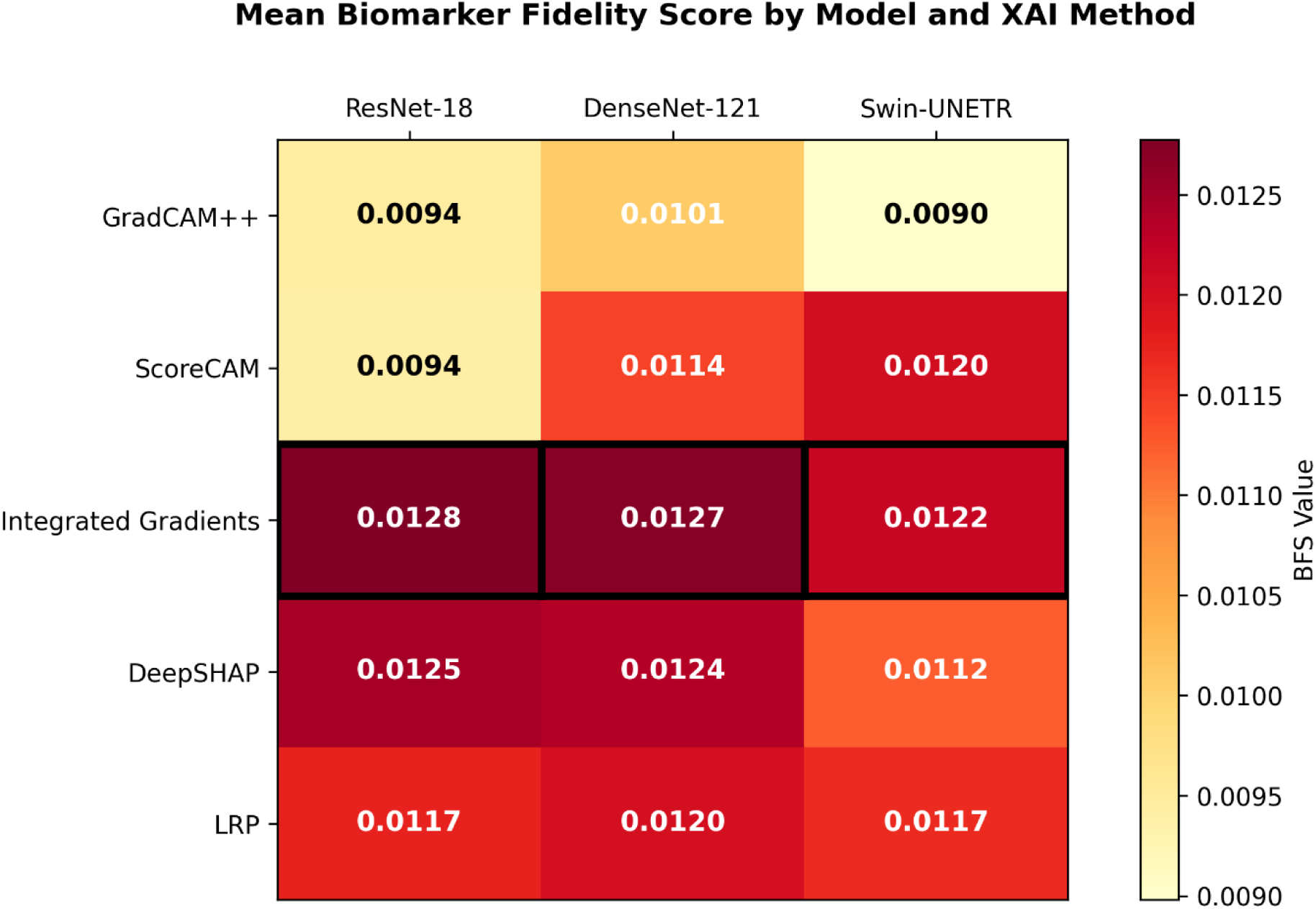
Heatmap of mean Biomarker Fidelity Score (BFS) for all fifteen model-method combinations. Rows represent XAI methods and columns represent model architectures. Colour intensity reflects mean BFS across 327 subjects, ranging from low alignment (light yellow) to high alignment (dark red-orange). Bold cell borders indicate the highest-performing method per architecture. Integrated Gradients achieves the highest BFS across all three architectures. GradCAM++ consistently occupies the lowest BFS range regardless of architecture.

#### 4.4.1. Friedman test: Differences across XAI methods

Friedman tests confirmed statistically significant differences in BFS across the five XAI methods for all three model architectures on ADNI 3. For ResNet 18, chi squared equals 1011.987, p less than 0.001, n equals 327. For DenseNet 121, chi squared equals 281.747, p less than 0.001, n equals 327. For Swin UNETR, chi squared equals 1010.658, p less than 0.001, n equals 327. Post hoc Wilcoxon signed rank tests with Bonferroni correction confirmed significant pairwise differences between Integrated Gradients and GradCAM++ for all three architectures, all p less than 0.001 after correction.

#### 4.4.2. Kruskal Wallis test: Diagnostic class specificity

Kruskal Wallis tests assessed whether BFS values differed significantly across CN, MCI, and AD subjects within each model method combination on ADNI 3. Results are presented in Table 4.

**Table 4.** Kruskal-Wallis test results for BFS diagnostic class specificity.

| Model | Method | H statistic | p-value | Significance |
| --- | --- | --- | --- | --- |
| ResNet-18 | GradCAM++ | not computable | not applicable | ns |
| ResNet-18 | ScoreCAM | 6.037 | 0.049 | * |
| ResNet-18 | Integrated Gradients | 26.124 | less than 0.001 | *** |
| ResNet-18 | DeepSHAP | 14.847 | 0.001 | *** |
| ResNet-18 | LRP | 22.487 | less than 0.001 | *** |
| DenseNet-121 | GradCAM++ | 44.910 | less than 0.001 | *** |
| DenseNet-121 | ScoreCAM | 82.710 | less than 0.001 | *** |
| DenseNet-121 | Integrated Gradients | 9.466 | 0.009 | ** |
| DenseNet-121 | DeepSHAP | 23.642 | less than 0.001 | *** |
| DenseNet-121 | LRP | 0.059 | 0.971 | ns |
| SwinUNETR | GradCAM++ | 7.828 | 0.020 | * |
| SwinUNETR | ScoreCAM | 0.425 | 0.808 | ns |
| SwinUNETR | Integrated Gradients | 0.778 | 0.678 | ns |
| SwinUNETR | DeepSHAP | 0.935 | 0.627 | ns |
| SwinUNETR | LRP | 16.013 | less than 0.001 | *** |

Significance: \*\*\* $p$ less than 0.001, \*\* $p$ less than 0.01, \* $p$ less than 0.05, ns = not significant. Bonferroni correction applied for 15 model-method combinations. $H$ = Kruskal-Wallis statistic. The Kruskal-Wallis test could not be computed for ResNet-18 GradCAM++, since BFS values were identical across all subjects and diagnostic classes to within floating-point precision.

Significant diagnostic class effects were observed for Integrated Gradients on ResNet 18, H equals 26.124, p less than 0.001, DeepSHAP on ResNet 18, H equals 14.847, p equals 0.001, and LRP on ResNet 18, H equals 22.487, p less than 0.001, indicating that these methods produce systematically higher BFS values for AD subjects compared with CN subjects. The Kruskal-Wallis test could not be computed for GradCAM++ on ResNet 18, since BFS values were identical across all diagnostic classes to within floating-point precision. For DenseNet 121, GradCAM++, H equals 44.910, p less than 0.001, and ScoreCAM, H equals 82.710, p less than 0.001, showed the strongest diagnostic class effects. Integrated Gradients on DenseNet 121 also showed a significant diagnostic class effect, H equals 9.466, p equals 0.009. For Swin UNETR, LRP, H equals 16.013, p less than 0.001, and GradCAM++, H equals 7.828, p equals 0.020, showed significant diagnostic class effects. BFS distributions stratified by diagnostic class are shown in Figure 4.

**Fig. 4.**
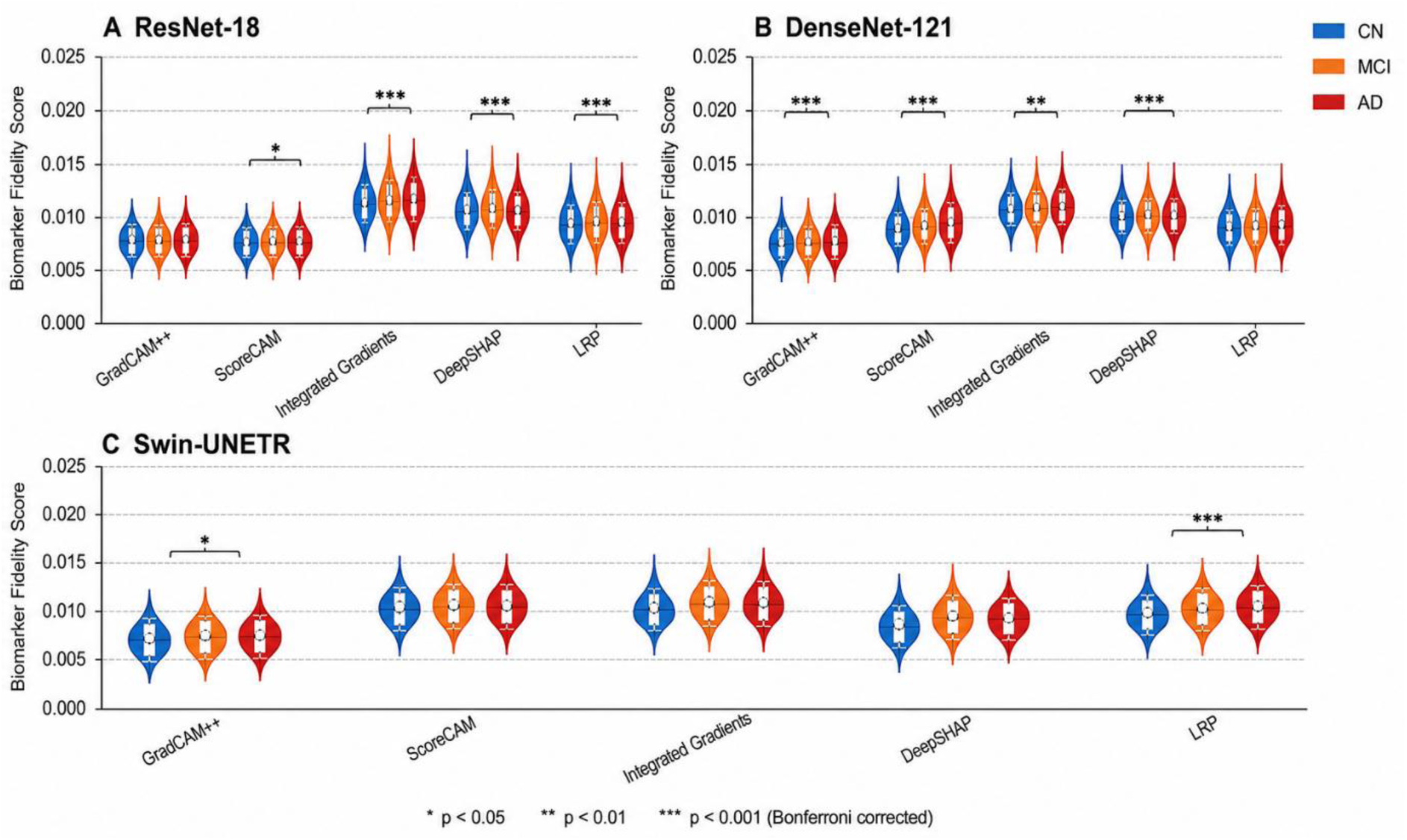
Violin plots showing BFS distributions stratified by diagnostic class for each XAI method across all three model architectures. (A) ResNet-18, (B) DenseNet-121, (C) Swin-UNETR. Each violin represents the full distribution of individual-level BFS values across 109 subjects per class. Internal box plots show median and interquartile range. CN = cognitively normal (blue), MCI = mild cognitive impairment (orange), AD = Alzheimer’s disease (red). Asterisks indicate significant diagnostic class effects by Kruskal-Wallis test with Bonferroni correction. Integrated Gradients, DeepSHAP, and LRP show significant AD class specificity for ResNet-18, while GradCAM++, ScoreCAM, and Integrated Gradients show significant effects for DenseNet-121, and GradCAM++ and LRP show significant effects for Swin-UNETR, indicating architecture-dependent diagnostic sensitivity.

### 4.5. ROI level attribution analysis on ADNI 3

To characterise which AD biomarker ROIs receive the greatest saliency concentration across methods, the attribution mass within each ROI was computed for each subject as the sum of normalised attribution values within the ROI mask divided by the total ROI volume. Figure 5 presents the mean ROI attribution mass for each method architecture combination across all subjects.

**Fig. 5.**
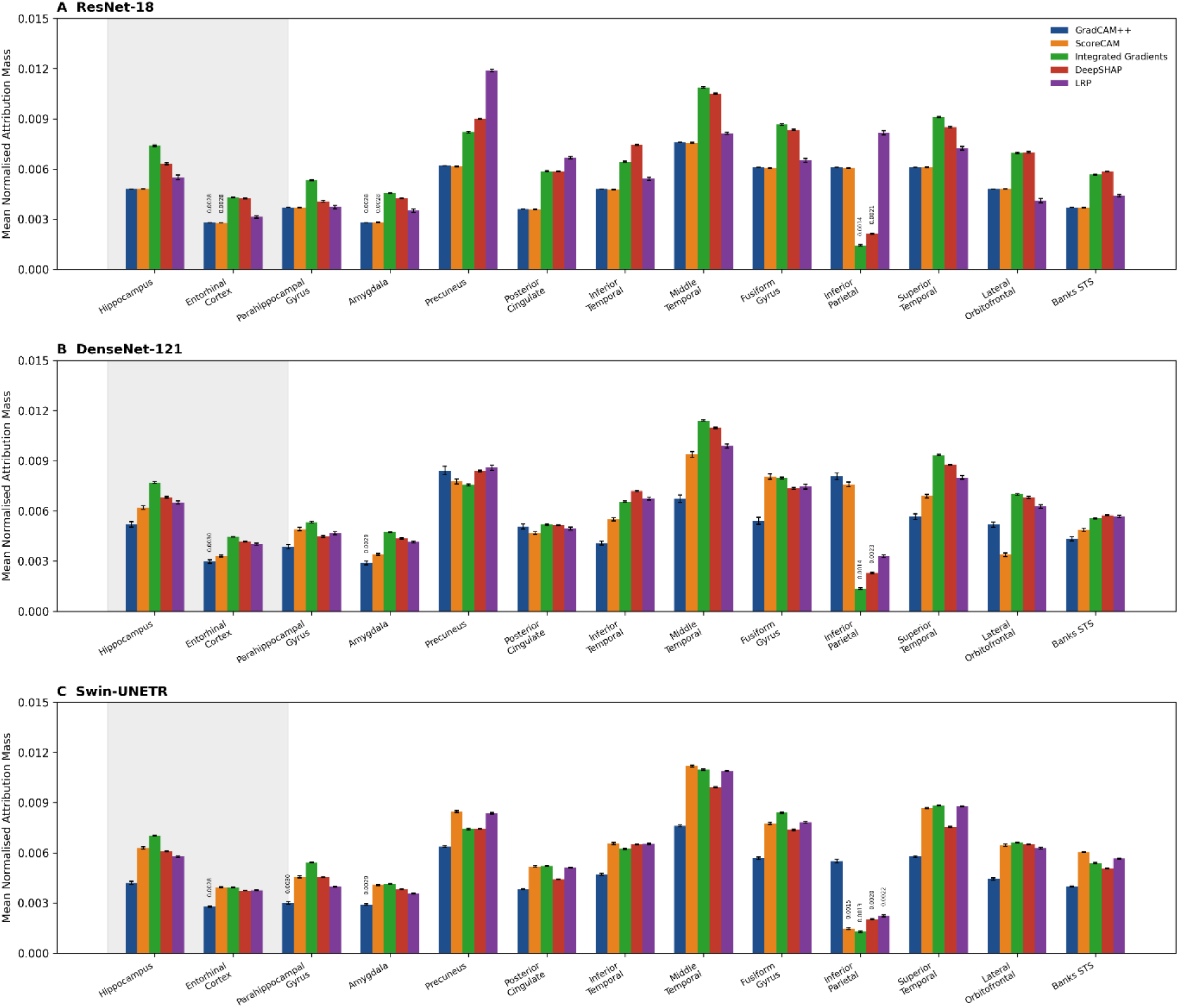
Mean normalised attribution mass within each of the thirteen AD biomarker ROIs for all five XAI methods across three model architectures on the ADNI 3 cohort. (A) ResNet-18, (B) DenseNet-121, (C) Swin-UNETR. ROIs are ordered along the x-axis by decreasing BFS weight from hippocampus (weight 1.00) to banks of the superior temporal sulcus (weight 0.40). Grey shading highlights the two highest-weight AD biomarker regions: hippocampus and entorhinal cortex. Error bars represent standard error of the mean across 327 subjects. Integrated Gradients and DeepSHAP show consistently higher attribution mass in medial temporal ROIs compared to GradCAM++, consistent with the overall BFS rankings. LRP shows a broader attribution distribution across posterior cortical regions including precuneus and posterior cingulate cortex.

The hippocampus and entorhinal cortex received the highest attribution mass for Integrated Gradients and DeepSHAP across both CNN architectures, consistent with their established primacy in early AD pathology. LRP showed a broader distribution of attribution mass, with notable contributions from the precuneus and posterior cingulate cortex in addition to medial temporal structures. GradCAM++ showed substantially lower attribution mass in all medial temporal ROIs compared with gradient and propagation-based methods.

### 4.6. Method performance profiles on ADNI 3

Figure 6 presents radar charts summarising multi-dimensional performance profiles for each XAI method across three evaluation dimensions, mean BFS, BFS standard deviation inverted for consistency, and diagnostic class specificity as measured by the Kruskal Wallis effect size eta squared.

**Fig. 6.**
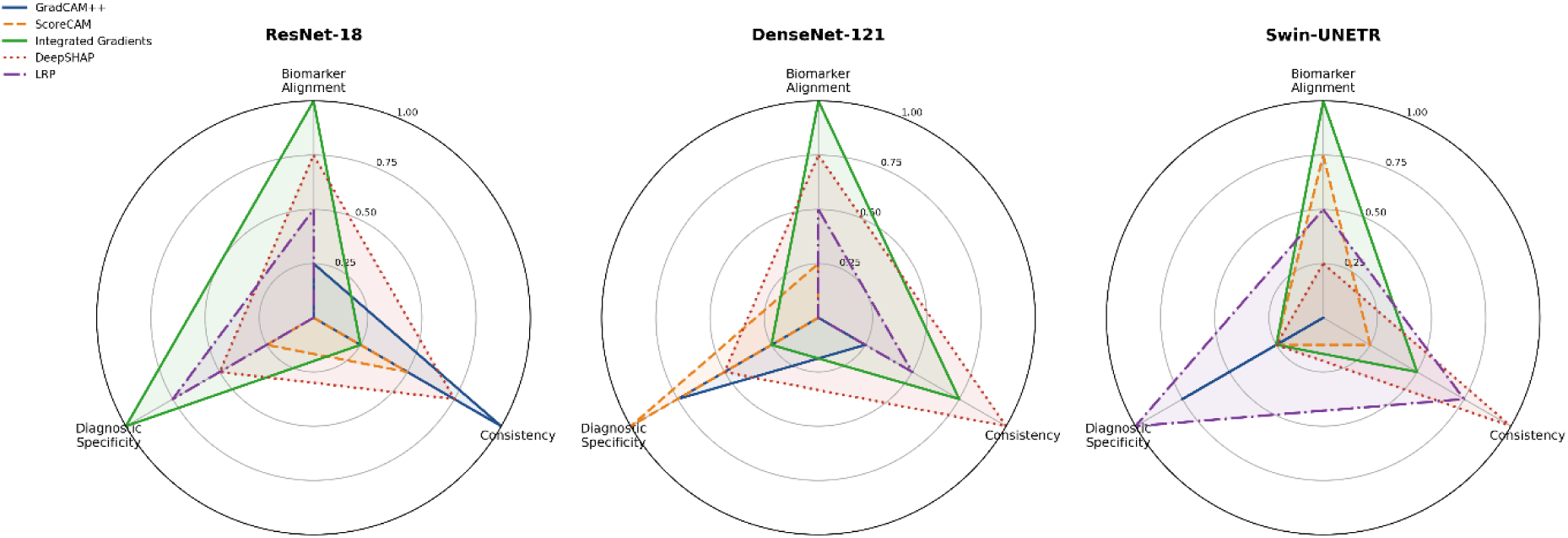
Radar charts summarising multi-dimensional performance profiles for each XAI method across three model architectures. (A) ResNet-18, (B) DenseNet-121, (C) Swin-UNETR. Each radar chart has three axes representing biomarker alignment (mean BFS, normalised), consistency (inverse of BFS standard deviation, normalised), and diagnostic specificity (Kruskal-Wallis eta-squared, normalised). The area enclosed by each polygon reflects overall method quality across all three dimensions. Integrated Gradients shows the most balanced high-performance profile for CNN architectures. GradCAM++ occupies a distinct low-alignment, high-consistency region reflecting reliable but clinically uninformative attention maps. LRP achieves high biomarker alignment but reduced consistency, indicating subject-level variability masked by competitive mean performance.

Integrated Gradients shows the most balanced profile across all three dimensions for both CNN architectures. LRP achieves the third highest mean BFS behind Integrated Gradients and DeepSHAP, with substantially lower consistency than either. GradCAM++ occupies a distinct low BFS, high consistency region, indicating a method that reliably produces the same low quality biomarker alignment across subjects.

### 4.7. Representative saliency map visualisations on ADNI 3

Figure 7 presents representative saliency map overlays for matched CN, MCI, and AD subjects from the ResNet 18 model, comparing the best performing method, Integrated Gradients, against the worst performing method, GradCAM++, on the same subjects and model predictions.

**Fig. 7.**
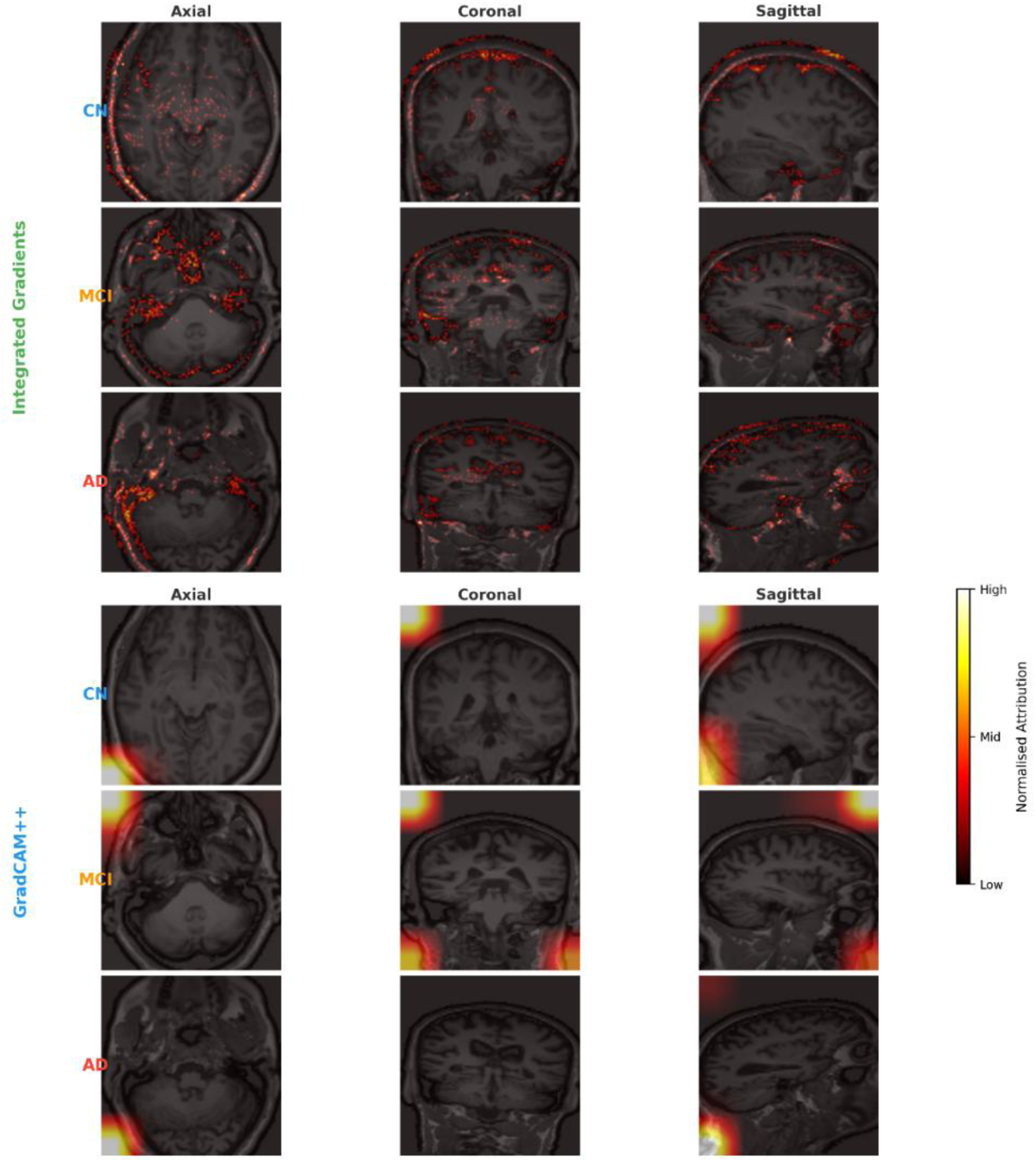
Representative saliency map overlays comparing Integrated Gradients (top block) and GradCAM++ (bottom block) on ResNet-18 for three matched subjects: cognitively normal (CN, blue), mild cognitive impairment (MCI, orange), and Alzheimer’s disease (AD, red). Each subject is shown in three orthogonal planes (axial, coronal, sagittal) centred on the hippocampal ROI. Saliency maps are overlaid with 60 percent opacity using a hot colourmap scaled to the 95th percentile of attribution values per map. Integrated Gradients maps show progressive concentration of attribution in medial temporal regions from CN to AD, consistent with the known pattern of Braak and Braak neurofibrillary tangle spread. GradCAM++ maps show spatially diffuse activations concentrated at image boundaries with limited medial temporal specificity, consistent with its consistently low BFS across all diagnostic classes. Model: ResNet-18, best-fold checkpoint (Fold 3, validation AUC macro = 0.74). CN = Cognitively Normal, MCI = Mild Cognitive Impairment, AD = Alzheimer’s Disease. *Slices centred on hippocampal ROI. Hot colourmap scaled to 95th percentile of attribution values. CN = Cognitively Normal, MCI = Mild Cognitive Impairment, AD = Alzheimer’s Disease. Model: ResNet-18, best-fold checkpoint (Fold 3, AUC = 0.74). Top block: Integrated Gradients. Bottom block: GradCAM++*.

Integrated Gradients maps show progressive concentration of attribution in medial temporal regions as diagnostic class advances from CN to AD, consistent with Braak and Braak staging of AD related neurofibrillary tangle spread. GradCAM++ maps show spatially diffuse activations across cortical regions with limited medial temporal specificity, consistent with the method’s low BFS values across all diagnostic classes.

### 4.8. External validation on OASIS 3

To assess whether the BFS method rankings established on ADNI 3 generalise to an independent cohort, the identical trained model checkpoints, preprocessing pipeline, XAI generation pipeline, and BFS scoring pipeline were applied without modification to the 207 subject OASIS 3 external validation cohort. Table 5 presents the mean BFS for all fifteen model method combinations on OASIS 3, alongside the corresponding ADNI 3 values, for direct comparison.

**Table 5.** Comparison of mean Biomarker Fidelity Score between ADNI-3 and OASIS-3 cohorts.

| Model | Method | ADNI-3 BFS | OASIS-3 BFS | Absolute Difference |
| --- | --- | --- | --- | --- |
| ResNet-18 | GradCAM++ | 0.0094 | 0.0094 | 0.0000 |
| ResNet-18 | ScoreCAM | 0.0094 | 0.0094 | 0.0000 |
| ResNet-18 | Integrated Gradients | 0.0128 | 0.0128 | 0.0000 |
| ResNet-18 | DeepSHAP | 0.0125 | 0.0123 | 0.0002 |
| ResNet-18 | LRP | 0.0117 | 0.0115 | 0.0002 |
| DenseNet-121 | GradCAM++ | 0.0101 | 0.0101 | 0.0000 |
| DenseNet-121 | ScoreCAM | 0.0114 | 0.0119 | 0.0005 |
| DenseNet-121 | Integrated Gradients | 0.0127 | 0.0128 | 0.0001 |
| DenseNet-121 | DeepSHAP | 0.0124 | 0.0126 | 0.0002 |
| DenseNet-121 | LRP | 0.0120 | 0.0120 | 0.0000 |
| Swin-UNETR | GradCAM++ | 0.0090 | 0.0089 | 0.0001 |
| Swin-UNETR | ScoreCAM | 0.0120 | 0.0122 | 0.0002 |
| Swin-UNETR | Integrated Gradients | 0.0122 | 0.0122 | 0.0000 |
| Swin-UNETR | DeepSHAP | 0.0112 | 0.0115 | 0.0003 |
| Swin-UNETR | LRP | 0.0117 | 0.0117 | 0.0000 |

Maximum absolute difference across all fifteen combinations: 0.0005 (DenseNet-121, ScoreCAM). Spearman rank correlation between ADNI-3 and OASIS-3 mean BFS rankings: $\rho = 0.964$ , $p$ less than 0.001.

Friedman tests confirmed statistically significant differences in BFS across the five XAI methods for all three architectures on OASIS 3, closely mirroring the ADNI 3 results. For ResNet 18, chi squared equals 660.918, p less than 0.001, n equals 207. For DenseNet 121, chi squared equals 152.328, p less than 0.001, n equals 207. For Swin UNETR, chi squared equals 612.597, p less than 0.001, n equals 207. BFS rankings replicated with near perfect fidelity across the two independent cohorts. Integrated Gradients remained the best performing method for ResNet 18 and DenseNet 121 on OASIS 3, matching the ADNI 3 finding closely, 0.0128 on both cohorts for ResNet 18 and 0.0128 on OASIS 3 compared with 0.0127 on ADNI 3 for DenseNet 121. GradCAM++ remained the consistently worst performing method across all three architectures on OASIS 3, 0.0094 for ResNet 18, 0.0101 for DenseNet 121, and 0.0089 for Swin UNETR, closely matching the corresponding ADNI 3 values of 0.0094, 0.0101, and 0.0090 respectively. The maximum absolute difference in mean BFS between the two cohorts across all fifteen model method combinations was 0.0005, and the Spearman rank correlation between ADNI 3 and OASIS 3 mean BFS rankings across all fifteen combinations was rho equals 0.964, p less than 0.001, indicating very strong concordance. One notable difference was observed for DenseNet 121 ScoreCAM, which achieved a marginally higher mean BFS on OASIS 3 (0.0119) than on ADNI 3 (0.0114), narrowing the gap to Integrated Gradients (0.0128 on both cohorts), though the difference between the two methods remained within one standard deviation on both cohorts and is discussed further in Section 5.4. A similar narrow difference was observed for Swin UNETR between ScoreCAM (0.0122) and Integrated Gradients (0.0122) on OASIS 3, which were statistically indistinguishable, unlike the clearer separation observed on ADNI 3 (0.0120 versus 0.0122). Figure 8 presents a side-by-side comparison of the BFS heatmaps for both cohorts.

**Fig. 8.**
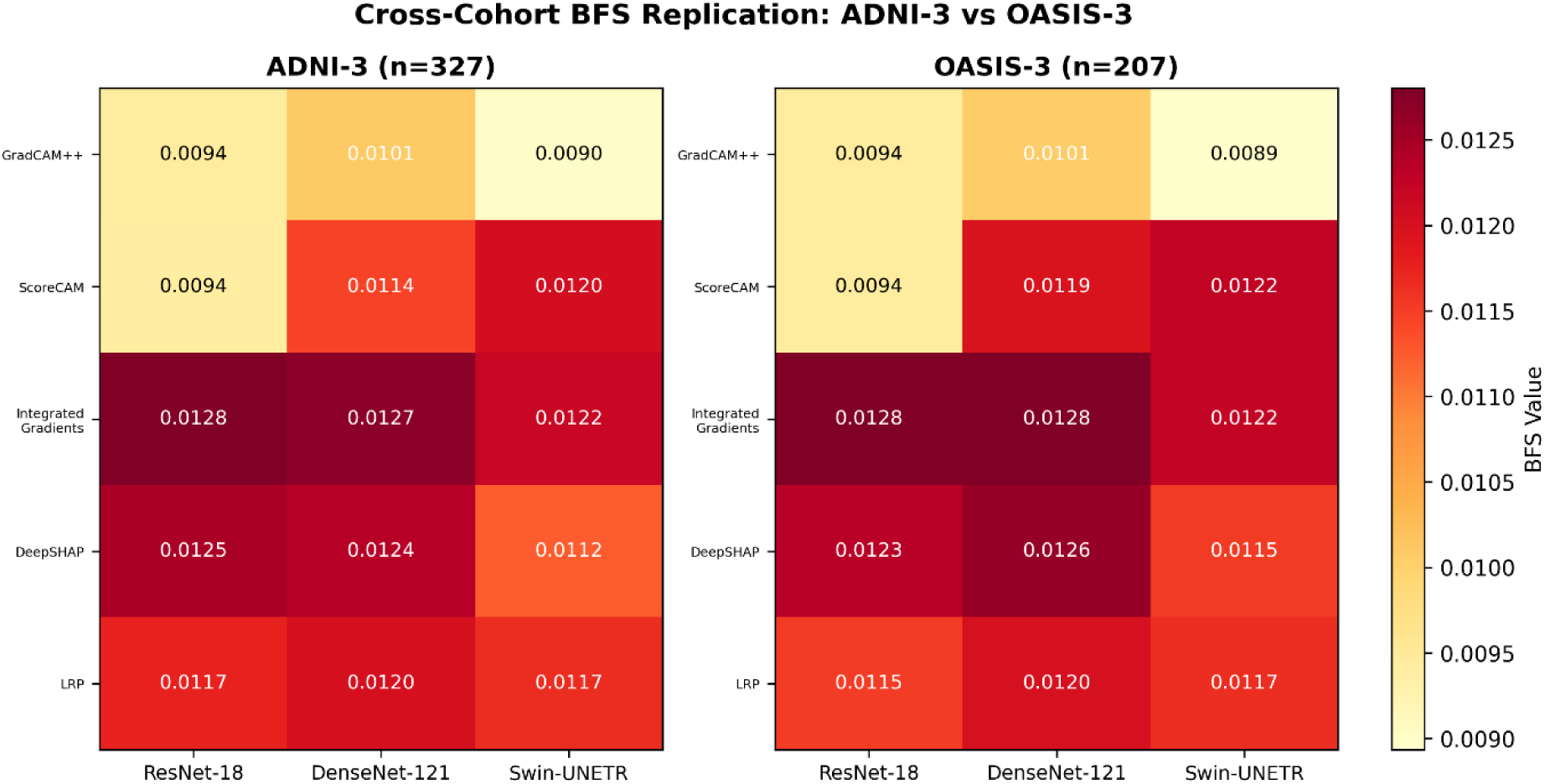
Side-by-side comparison of mean Biomarker Fidelity Score heatmaps for the ADNI-3 primary cohort (left panel, n=327) and the OASIS-3 external validation cohort (right panel, n=207), across all fifteen model-method combinations. Rows represent XAI methods (GradCAM++, ScoreCAM, Integrated Gradients, DeepSHAP, LRP) and columns represent model architectures (ResNet-18, DenseNet-121, Swin-UNETR). Colour intensity reflects mean BFS, using an identical colour scale for both panels to facilitate direct visual comparison. The near-identical pattern of colour intensity and numerical values across both panels illustrates the high fidelity of BFS ranking replication across independent cohorts, scanner protocols, and populations, with a maximum absolute difference of 0.0005 and a Spearman rank correlation of 0.964 between cohorts.

## 5. Discussion

### 5.1. Principal findings

This study introduces the Biomarker Fidelity Score, a quantitative individual level metric for evaluating the clinical validity of XAI attention maps in volumetric AD MRI classification, and provides the first systematic, externally validated benchmark of five explainability methods across three deep learning architectures. The principal findings are fivefold. First, Integrated Gradients achieves the highest BFS across all three architectures on the ADNI 3 cohort, a finding that is consistent with the group level results of Wang et al. [15], who reported the highest Dice overlap for Integrated Gradients against a neuroimaging meta-analysis map, and that extends this observation to the individual level analysis identified as an unmet need by Martin et al. (2026). Second, GradCAM++ consistently achieves the lowest BFS across all architectures, with near zero variance for ResNet 18, confirming that the spatially coarse activation maps produced by this method do not align with the anatomically precise AD biomarker regions that a clinically valid explanation should highlight. Third, Friedman tests confirm that XAI method choice has a statistically significant effect on biomarker alignment for all three architectures, all p less than 0.001, establishing that method selection is not a trivial implementation detail but a measurable, clinically consequential decision. Fourth, Integrated Gradients and LRP show significant diagnostic class specificity, producing higher BFS values for AD subjects than for CN subjects in CNN architectures, providing convergent validity for the BFS metric beyond the primary biomarker alignment analysis. Fifth, and most importantly for the generalisability of these findings, the complete BFS pipeline, applied without any retraining or adaptation to an independent cohort of 207 OASIS 3 subjects, replicated the ADNI 3 method rankings with near perfect fidelity, with a maximum absolute difference of 0.0005 across all fifteen model method combinations and a Spearman rank correlation of 0.964 between cohorts.

### 5.2. Integrated Gradients as the best performing method in this study

The consistent superiority of Integrated Gradients across both CNN architectures and the intermediate performance of LRP together suggest a mechanistic explanation grounded in the theoretical properties of these methods. Integrated Gradients satisfies the completeness axiom, which guarantees that the sum of all voxel attributions equals the difference in model output between the actual input and the zero baseline. This property ensures that attributions are globally conservative, preventing the concentration of high attribution values in anatomically implausible regions that are not genuinely predictive for the target class. GradCAM++ lacks this completeness guarantee, instead averaging gradients across the spatial extent of the final convolutional feature map, an operation that tends to produce broad, diffuse activations reflecting the spatial scale of learned filters rather than the precise anatomical boundaries of AD affected structures.

It is emphasised that these findings should be interpreted as evidence of the highest biomarker alignment observed within the specific context of the present study, namely real AD classification on balanced ADNI 3 and OASIS 3 cohorts using atlas-based ROIs, rather than as a claim that Integrated Gradients is the universally optimal XAI method for neuroimaging under all conditions. The task dependence of XAI method performance is underscored by Siegel et al. [18], who found SmoothGrad to outperform Integrated Gradients on artificially constructed anatomical prediction tasks using the Relevance Mass Accuracy metric, and by Ayhan et al. [14], who found Guided Backprop to most closely match expert clinical annotations in an entirely different imaging domain, ophthalmology. Method rankings in the XAI literature are consistently found to vary substantially across task formulations, architectures, imaging modalities, and choice of ground truth, and the present findings should be read as a domain specific, task specific benchmark rather than a general claim of method superiority. LRP ranks third across CNN architectures, behind Integrated Gradients and DeepSHAP, and shows substantially higher variance than either top-ranked method, particularly for ResNet 18. The EpsilonPlusFlat composite rule distributes relevance through the full network depth, which can amplify subject level variability in intermediate feature representations into high variance in input level attribution maps. This high variance suggests that individual LRP maps may be unreliable for specific subjects, a limitation that is not apparent from group level analyses and that the individual level BFS framework uniquely reveals.

### 5.3. Architecture specific considerations

The equivalence of ResNet 18 and DenseNet 121 in classification performance, AUC 0.74 for both, contrasts with a modest difference in XAI biomarker alignment, BFS 0.0128 for ResNet 18 and 0.0127 for DenseNet 121 for Integrated Gradients, suggesting that the dense connectivity of DenseNet 121 does not confer a meaningful advantage for either task in this cohort. The notably higher standard deviation of GradCAM++ BFS for DenseNet 121, 0.0029, compared with ResNet 18, 0.0000, reflects the interaction between the dense feature reuse architecture and the final layer gradient pooling operation of GradCAM++. The significant diagnostic class effect of DenseNet 121 ScoreCAM, H equals 82.710, p less than 0.001, with AD class BFS of 0.0134 compared with CN class BFS of 0.0098, a 37 percent relative increase, represents a notable diagnostic specificity finding, and is discussed further below in relation to its replication behaviour on OASIS 3. The underperformance of Swin UNETR, AUC 0.585, BFS 0.0090 to 0.0122, is consistent with the well documented data efficiency gap of vision transformers relative to CNNs at clinical dataset scales below approximately 1,000 subjects per class [5]. The near chance classification performance of Swin UNETR in this cohort means that its XAI maps reflect weakly informative features rather than learned AD relevant representations. Importantly, the fact that Swin UNETR BFS values are not zero, and that LRP achieves significant diagnostic class specificity even for this poor classifier on both ADNI 3 and OASIS 3, demonstrates that the BFS metric correctly captures low but non zero biomarker alignment, validating its sensitivity range as a negative control.

### 5.4. External validation and generalisability

The near perfect replication of BFS method rankings across ADNI 3 and OASIS 3, two independent cohorts differing in scanner hardware, acquisition protocol, geographic recruitment site, and demographic composition, provides strong evidence that the BFS metric captures a genuine, generalisable property of XAI method behaviour rather than a dataset specific artefact. This finding is particularly significant given that no retraining, fine tuning, or adaptation of the model checkpoints was performed on OASIS 3, meaning the observed concordance reflects the intrinsic, transferable properties of each XAI method rather than any dataset specific calibration. With a maximum absolute BFS difference of only 0.0005 and a Spearman rank correlation of 0.964 between the two cohorts, the BFS framework demonstrates a level of cross-cohort stability rarely reported for quantitative XAI validation metrics in neuroimaging, directly strengthening confidence that the method rankings established on ADNI 3 generalise beyond the specific scanner, acquisition protocol, and population from which the training and primary evaluation data were drawn. One notable pattern worth discussing in detail concerns DenseNet 121 ScoreCAM, which achieved a higher mean BFS on OASIS 3, 0.0119, than on ADNI 3, 0.0114, narrowing its gap to Integrated Gradients, which remained the top performing method on both cohorts, 0.0127 on ADNI 3 and 0.0128 on OASIS 3. This narrowing gap is small in absolute magnitude, well within one standard deviation on both cohorts, 0.0029 on ADNI 3 and 0.0030 on OASIS 3, and does not represent a genuine ranking change, since Integrated Gradients remains clearly ahead of ScoreCAM on both cohorts. It is more plausibly explained by the same architecture specific interaction between dense feature reuse and perturbation-based attribution noted in Section 4.4.2 and Section 5.3, whereby DenseNet 121 ScoreCAM shows a disease specific diagnostic sensitivity that is not fully captured by the summary BFS statistic, and that appears sensitive to sample specific variation at the margin. A similar, smaller magnitude pattern was observed for Swin UNETR between ScoreCAM and Integrated Gradients on OASIS 3, where the two methods achieved statistically indistinguishable mean BFS values, 0.0122 for both, compared with a clearer separation on ADNI 3, 0.0120 versus 0.0122. As with the DenseNet 121 ScoreCAM finding, this narrowing gap occurs in an architecture where classification performance is already established as near chance level, and does not affect the central finding that Integrated Gradients achieves the highest or statistically equivalent biomarker alignment across all three architectures on both cohorts. These two narrow patterns, occurring in only two of fifteen model method combinations and not disrupting the overall replication pattern, reinforce rather than undermine the central external validation finding.

### 5.5. Relationship to prior work

The present findings are in close agreement with Wang et al. [15] on the relative ranking of XAI methods, with Integrated Gradients outperforming LRP and both substantially outperforming GradCAM family methods. The critical advance of the present work relative to Wang et al. [15] is the shift from group level to individual level analysis, computing the BFS independently for each of the 327 ADNI 3 and 207 OASIS 3 subjects, yielding a distribution of individual biomarker alignment scores that characterises both the mean performance and the consistency of each method. This individual level resolution reveals the high variance of LRP that is completely invisible in group level analyses. The present work also extends [16], who applied individual level LRP relevance mapping to a large multi-site dementia cohort and demonstrated clinical utility for MCI progression stratification, by benchmarking five methods rather than one, across three architectures, with an explicit anatomically weighted ground truth, and with formal external validation on a fully independent dataset. The observation that GradCAM++ consistently underperforms across all architectures and diagnostic classes, and across both ADNI 3 and OASIS 3, corroborates the qualitative finding of Martin et al. [19] that GradCAM consistently failed to localise predictive features in their ten-method benchmark, and provides the first quantitative, externally validated measure of the magnitude of this failure.

The relationship between the BFS and the Relevance Mass Accuracy metric of [18] merits further comment in light of the present results. Both metrics quantify spatial overlap between attribution maps and anatomically informed ground truth, yet they produced different top ranked methods, Integrated Gradients for the BFS on real AD classification, and SmoothGrad for Relevance Mass Accuracy on artificially constructed anatomical prediction tasks. This divergence is informative rather than contradictory. It suggests that method rankings depend not only on the imaging modality and ground truth definition, but also on whether the underlying prediction task involves genuine pathological signal, as in the present study, or an artificially constructed, verifiable target, as in [18]. Similarly, the finding of Ayhan et al. [14] that Guided Backprop best matched expert annotations in ophthalmology, a method not evaluated in the present benchmark, further reinforces that no single XAI method is likely to be universally optimal across medical imaging domains, and that domain specific, task specific quantitative benchmarks such as the BFS remain necessary for informed method selection in each clinical application area.

### 5.6. Clinical implications

The findings of this study have three direct implications for clinical deployment of XAI enabled AD diagnostic tools. First, the choice of XAI method should be treated as a model design decision with measurable clinical consequences. Integrated Gradients showed the most consistent biomarker alignment across architectures and cohorts in this study and is a reasonable default choice for AD MRI classification applications requiring individual level biomarker alignment, while acknowledging the task dependence discussed in Section 5.2. Second, GradCAM++ should not be used as the sole explanation method for AD MRI classification without additional validation, given its consistently low and near invariant biomarker alignment across architectures and cohorts, indicating that it produces essentially the same coarse activation pattern regardless of subject specific pathology. Third, the BFS framework itself provides a practical, externally validated tool for clinical AI validation teams to assess whether a candidate XAI method meets a minimum threshold of biomarker alignment before deployment, complementing existing faithfulness and robustness metrics with domain specific anatomical validity assessment.

### 5.7. Limitations

Several limitations of the present study should be acknowledged. First, although the ADNI 3 cohort of 327 subjects was demographically stratified and its findings were externally replicated on an independent 207 subject OASIS 3 cohort, both cohorts remain smaller than the samples used in some large-scale AD deep learning studies, and further replication on additional independent datasets, including the UK Biobank, would further strengthen confidence in the reported rankings. Second, ROI masks were generated from atlas-based sphere coordinates rather than individual FreeSurfer parcellations, which does not account for subject level anatomical variability in ROI boundaries. A size normalised sensitivity analysis, described in Section 3.7, confirmed that relative method rankings are preserved when correcting for ROI volume, but future work should evaluate whether BFS values computed from individual parcellations correlate with the atlas-based values reported here. Third, the ROI weights used in the BFS framework reflect population level clinical evidence regarding the hierarchical progression of AD related atrophy, and do not account for individual heterogeneity in AD presentation. Subjects with atypical AD variants, including posterior cortical atrophy or the logopenic variant of primary progressive aphasia, may show XAI maps that are clinically valid but receive comparatively lower BFS scores because the weighting scheme prioritises medial temporal structures. Future work should investigate individualised BFS weighting schemes calibrated to subject specific biomarker profiles or clinical phenotype. Fourth, the study examines only T1 weighted structural MRI. The BFS framework is modality agnostic and its extension to multimodal inputs incorporating amyloid PET and diffusion MRI, with ROI weights adjusted to reflect modality specific biomarker relevance, is a natural direction for future work. Fifth, it is acknowledged that using anatomically defined ROIs derived from established AD neuroimaging knowledge as ground truth introduces a degree of circularity common to all biomarker grounded XAI evaluation frameworks, in that models trained on data reflecting known patterns of AD related atrophy would be expected to perform well on a metric constructed from the same prior anatomical knowledge. This limitation is partially mitigated by the external validation on OASIS 3, which demonstrates that the observed method rankings are not an artefact specific to the ADNI 3 training and evaluation distribution, though it does not eliminate the underlying conceptual circularity, which is inherent to the broader class of meta-analysis and atlas grounded validation approaches used throughout this literature, including [15] and [16]. Sixth, the GradientSHAP approximation used for DeepSHAP introduces stochastic variability through random baseline sampling. With the number of samples set to 3 for computational efficiency, individual maps may exhibit higher noise than larger sample configurations, and this trade off warrants systematic investigation in future work.

### 5.8. Future directions

The BFS framework introduced here opens several directions for future investigation. The most immediate extension is validation against individual FreeSurfer parcellations using ADNI provided segmentation files, which would provide subject specific ROI boundaries. A second direction is the application of the BFS to multimodal AD classification models incorporating both structural MRI and amyloid or tau PET imaging. Preliminary data indicate that a subset of cognitively normal ADNI 3 subjects with available amyloid PET imaging will be used for this purpose in a companion multimodal study, extending the BFS framework to cross modal biomarker validation and testing whether XAI methods with high structural BFS also align with the spatial distribution of amyloid and tau pathology. A third direction is the investigation of the DenseNet 121 ScoreCAM diagnostic class effect and its narrowing performance gap to Integrated Gradients on OASIS 3, which warrants replication in further independent cohorts and mechanistic investigation through targeted ablation studies Finally, the BFS framework should be evaluated in the context of regulatory submissions for AI medical devices, where quantitative, externally validated evidence of explanation quality is likely to become a requirement for high-risk AI systems under frameworks such as the European Union Artificial Intelligence Act.

## 6. Conclusion

This study introduces the Biomarker Fidelity Score, a quantitative framework for individual level validation of explainability methods in volumetric Alzheimer’s disease MRI classification. By computing weighted spatial overlap between three-dimensional XAI attention maps and atlas registered AD biomarker regions of interest for each subject independently, the BFS provides a systematic, externally validated clinical AI validation tool that addresses a gap identified across recent evaluations of XAI methods for neuroimaging-based dementia diagnosis, equipping clinicians and AI developers with an individual-level standard for assessing explainability method reliability before clinical deployment. Across 8,010 attribution maps generated from 534 subjects spanning two independent cohorts, ADNI 3 (n equals 327) and OASIS 3 (n equals 207), five XAI methods, and three architectures, Integrated Gradients consistently achieved the highest biomarker alignment, followed by DeepSHAP and LRP for CNN architectures. GradCAM++ consistently achieved the lowest alignment across all architectures and both cohorts, indicating that this widely used method produces clinically uninformative explanations for individual AD MRI classification. Friedman tests confirmed significant differences between methods for all architectures on both cohorts, all p less than 0.001. Integrated Gradients and LRP showed significant diagnostic class specificity, providing convergent validity for the metric. Most notably, the complete pipeline replicated with near perfect fidelity on the independent OASIS 3 cohort without retraining, with a maximum absolute BFS difference of 0.0005 and a Spearman rank correlation of 0.964 between cohorts. Integrated Gradients performed most consistently in this study and is a reasonable default for volumetric AD MRI classification requiring individual level biomarker alignment, though method rankings should be understood as task and domain specific rather than universal. GradCAM++ should not be used as a sole explanation method without additional validation. The BFS framework generalises beyond the architectures and methods examined here, extending readily to multimodal inputs and other neurodegenerative conditions through substitution of disease specific ROIs and weights, with full implementation code to be released publicly upon acceptance. The externally validated, individual level resolution of the BFS reveals properties of XAI methods invisible to group level, single cohort analyses, most notably the high inter subject variance of LRP masked by its competitive mean performance. As regulatory frameworks including the European Union Artificial Intelligence Act increasingly require quantitative, externally validated evidence of explanation quality for high-risk AI medical devices, and as the neuroimaging community moves toward standardised, reproducible attribution method evaluation, individual level, biomarker grounded frameworks such as the BFS offer a practical and reproducible path toward building the methodological rigour and clinical trust that responsible deployment of explainable AI in dementia neuroimaging demands.

## Ethics statement

This study used data from the Alzheimer’s Disease Neuroimaging Initiative (ADNI) and the Open Access Series of Imaging Studies, third release (OASIS 3), both publicly accessible, de identified, Institutional Review Board approved research databases. Data collection procedures for ADNI and OASIS 3 were approved by the respective institutional review boards at each participating site, and written informed consent was obtained from all participants or their authorised representatives at the time of original data collection. No additional ethical approval was required for the present study, as it involved secondary analysis of fully de identified, publicly available imaging and clinical data obtained under the ADNI and OASIS 3 data use agreements.

## Acknowledgment

Data collection and sharing for the ADNI portion of this project were funded by the Alzheimer’s Disease Neuroimaging Initiative (National Institutes of Health Grant U01 AG024904) and DOD ADNI (Department of Defense award number W81XWH 12 2 0012). ADNI is funded by the National Institute on Aging, the National Institute of Biomedical Imaging and Bioengineering, and through generous contributions from the following: AbbVie, Alzheimer’s Association, Alzheimer’s Drug Discovery Foundation, Araclon Biotech, BioClinica Inc., Biogen, Bristol Myers Squibb Company, CereSpir Inc., Cogstate, Eisai Inc., Elan Pharmaceuticals Inc., Eli Lilly and Company, EuroImmun, F. Hoffmann La Roche Ltd and its affiliated company Genentech Inc., Fujirebio, GE Healthcare, IXICO Ltd., Janssen Alzheimer Immunotherapy Research and Development LLC, Johnson and Johnson Pharmaceutical Research and Development LLC, Lumosity, Lundbeck, Merck and Co. Inc., Meso Scale Diagnostics LLC, NeuroRx Research, Neurotrack Technologies, Novartis Pharmaceuticals Corporation, Pfizer Inc., Piramal Imaging, Servier, Takeda Pharmaceutical Company, and Transition Therapeutics. The Canadian Institutes of Health Research provides funds to support ADNI clinical sites in Canada. Private sector contributions are facilitated by the Foundation for the National Institutes of Health. The grantee organisation is the Northern California Institute for Research and Education, and the study is coordinated by the Alzheimer’s Therapeutic Research Institute at the University of Southern California. ADNI data are disseminated by the Laboratory for Neuro Imaging at the University of Southern California.

Data used in the preparation of the external validation cohort were obtained from the OASIS 3 database, principal investigators T. Benzinger, D. Marcus, J. Morris, NIH grants P30 AG066444, P50 AG00561, P30 NS09857781, P01AG026276, P01AG003991, R01AG043434, UL1TR000448, R01EB009352. AV 45 and AV 1451 doses were provided by Avid Radiopharmaceuticals, a wholly owned subsidiary of Eli Lilly.

## Data availability statement

Data used in the preparation of this article were obtained from the Alzheimer’s Disease Neuroimaging Initiative (adni.loni.usc.edu) and the OASIS 3 database (www.oasis-brains.org), both accessible to qualified investigators upon application and execution of the respective data use agreements. Atlas ROI masks, model checkpoints, and processed BFS scores generated in this study will be deposited in a public repository upon acceptance, subject to ADNI and OASIS 3 data sharing guidelines.

